# Coordination of host and endosymbiont gene expression governs endosymbiont growth and elimination in the cereal weevil *Sitophilus* spp

**DOI:** 10.1101/2023.04.03.535335

**Authors:** Mariana Galvão Ferrarini, Agnès Vallier, Carole Vincent-Monégat, Elisa Dell’Aglio, Benjamin Gillet, Sandrine Hughes, Ophélie Hurtado, Guy Condemine, Anna Zaidman-Rémy, Rita Rebollo, Nicolas Parisot, Abdelaziz Heddi

**Affiliations:** Univ Lyon, INSA Lyon, INRAE, BF2I, UMR 203, 69621 Villeurbanne, France; Université de Lyon, Université Lyon 1, CNRS, Laboratoire de Biométrie et Biologie Evolutive UMR 5558, F-69622 Villeurbanne, France; Univ Lyon, INRAE, INSA Lyon, BF2I, UMR 203, 69621 Villeurbanne, France; Institut de Génomique Fonctionnelle de Lyon (IGFL), CNRS UMR 5242, Ecole Normale Supérieure de Lyon, Université de Lyon, Lyon, France; Univ Lyon, Université Lyon 1, INSA de Lyon, CNRS UMR 5240 Microbiologie Adaptation et Pathogénie, Villeurbanne, France

## Abstract

**Background:** Insects living in nutritionally poor environments often establish long-term relationships with intracellular bacteria that supplement their diets and improve their adaptive and invasive powers. Even though these symbiotic associations have been extensively studied on physiological, ecological and evolutionary levels, few studies have focused on the molecular dialogue between host and endosymbionts to identify genes and pathways involved in endosymbiosis control and dynamics throughout host development.

**Results:** We simultaneously analyzed host and endosymbiont gene expression during the life cycle of the cereal weevil *Sitophilus oryzae*, from larval stages to adults, with a particular emphasis on emerging adults where the endosymbiont *Sodalis pierantonius* experiences a contrasted growth-climax-elimination dynamics. We unraveled a constant arms race in which different biological functions are intertwined and coregulated across both partners. These include immunity, metabolism, metal control, apoptosis, and bacterial stress response.

**Conclusions:** The study of these tightly regulated functions, which are at the center of symbiotic regulations, provides evidence on how hosts and bacteria finely tune their gene expression and respond to different physiological challenges constrained by insect development in a nutritionally limited ecological niche.

**Graphical Abstract:** 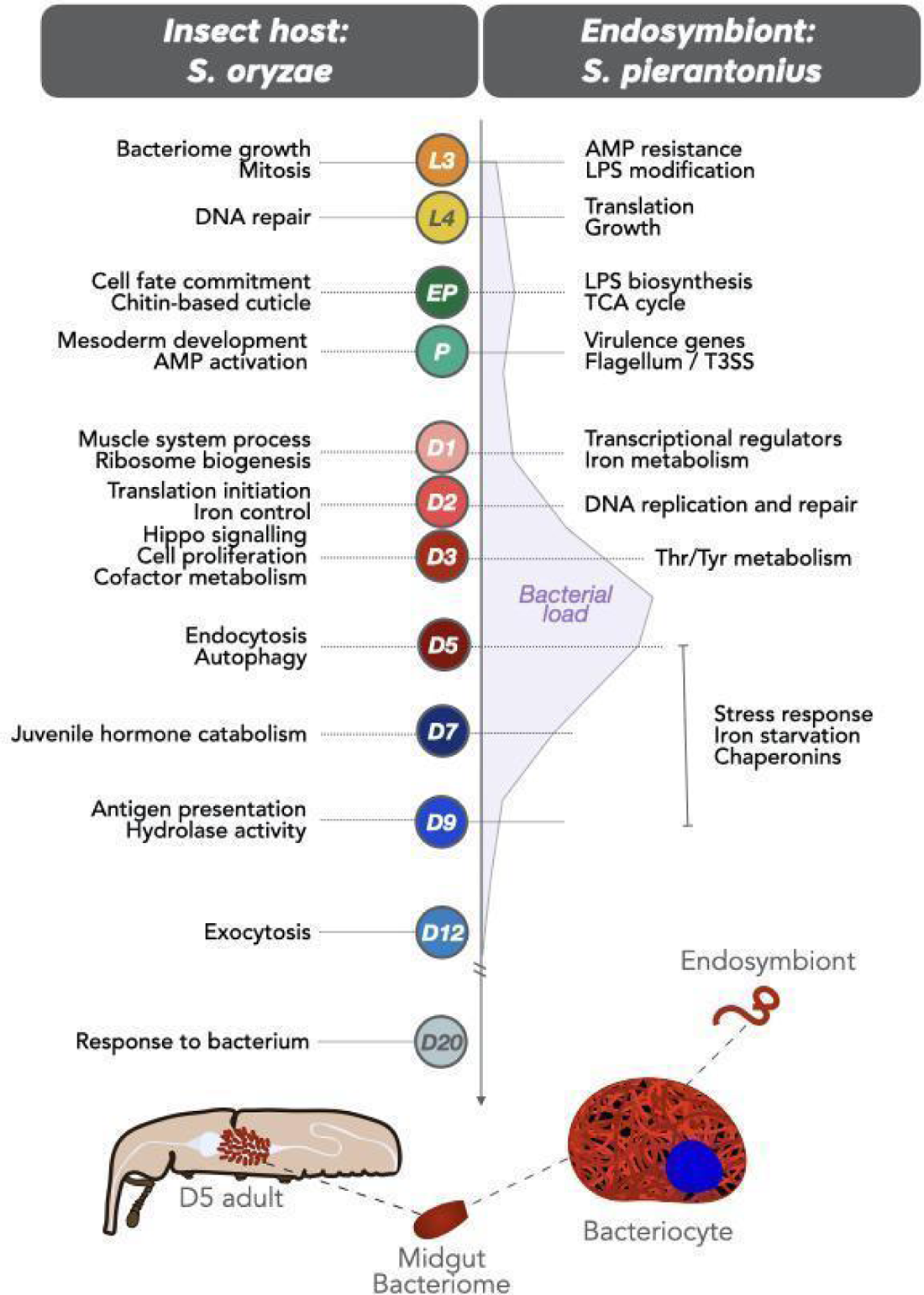

## Introduction

One peculiar attribute of many insects thriving on nutritionally unbalanced niches, including insect pests and disease vectors, is their ability to establish long-term relationships with heritable intracellular bacteria[1,2]. These so-called endosymbionts supplement hosts with metabolic components lacking in their diets and are known to impact organisms in terms of genome evolution, reproduction, host ecology, metabolic exchanges, and immune functions[3–10]. In many species, endosymbiont distribution is restricted to female germ cells and bacteriocytes, which are specialized cells that can group into a bacteriome organ that secludes the bacteria and prevents their exposure to the host immune system[11–13]. These intimate relationships likely emerged from an ancestral infection of free-living bacteria whose genomes experienced drastic size reduction, global enrichment in AT nucleic bases, and gene pseudogenization and deletion[14–20]. Moreover, vertically transmitted endosymbionts are under a relaxed evolutionary pressure due to a stable, nutritive and competition-free environment allowing for the accumulation of bacterial genome rearrangements and favoring host-symbiont co-evolution. This is highlighted by the loss of bacterial genes that become redundant or unnecessary in the new association, including virulence genes and genes encoding the bacterial cell wall elements[21,22], by the loss or erosion of most transcriptional regulatory elements[23,24], and by the increased metabolic interaction and complementation between the host and the endosymbiont[1,21,25–27].

Even though the metabolic, ecological and evolutionary aspects of endosymbiosis have been well studied, the molecular mechanisms underlying the maintenance of endosymbionts and their regulation concerning the host developmental needs, especially regarding the trade-off between bacterial burden and bacterial benefits, remain largely unexplored. Several studies focused on the model system Aphid-*Buchnera* and a recent work highlighted the coordination of host and endosymbiont genes in relation to bacterial titer[28]. Nevertheless, given the extensively reduced genome of *Buchnera aphidicola*, global transcriptomic analyses revealed weak gene expression changes in the bacteria[29,30] suggesting that endosymbiont regulation depends largely on the aphid host[31–34]. Thus, a comprehensive study on the regulation of host-endosymbiont interactions at critical stages of host development requires a one-to-one model system in which both the host and the bacteria are able to respond to different developmental constraints and biological stimuli with coordinated gene expression.

Endosymbiosis between the cereal weevil *Sitophilus oryzae* and the Gram-negative bacteria *Sodalis pierantonius* is most likely an exception among insects regarding several features. Contrary to their relatives, cereal weevils have established a symbiosis with *S. pierantonius* quite recently (<0.03 Myr)[35]. Despite having lost many genes compared to free-living relatives, the genome of *S. pierantonius* has not undergone a massive erosion, as observed in more ancient endosymbiotic bacteria, and encodes over 150 full-length regulatory proteins[19] as well as functional virulence factors[36,37]. In addition, the study of cereal weevils has the advantage of being associated solely with the *S. pierantonius* bacterium: most laboratory strains are free of *Wolbachia*, a common intracellular bacterium found in invertebrates[38], and remarkably, no commensal bacteria have been identified within their guts so far. While *S. pierantonius* has never been cultivated successfully *in vitro*, weevils can be artificially depleted of the bacteria (aposymbiotic insects)[39], allowing for comparative studies between symbiotic and aposymbiotic insects with the same genetic background. Finally, the cell dynamics of *S. pierantonius* undergoes a remarkably contrasted change during host development. Larval stages are considered homeostatic as the endosymbiont population grows slightly within a unique bacteriome, and bacteriocytes do not undergo any cellular or morphological changes. During metamorphosis, bacteriocytes move along the midgut and endosymbionts infect stem cells that differentiate into new bacteriocytes forming new bacteriomes at the apex of gut caeca[36]. In emerging adults, the endosymbiont population drastically increases (∼5 fold in five days)[40,41], which was shown to be associated with metabolic requirements of the host while building its cuticle. Once the cuticle is achieved, the endosymbionts are eliminated and completely recycled in 5 to 7 days. The midguts of young adults will remain devoid of bacteria throughout the rest of their lives. We have recently demonstrated there is a cost associated with such a drastic increase in bacterial load, suggesting a balance between the metabolic requirements associated with cuticle synthesis and the cost of harboring a large bacterial population[41]. Here, we focus on understanding the shift in endosymbiont control associated with the increase in bacterial load and the molecular signals directing endosymbiont elimination.

To demonstrate whether host and symbiont gene expression are coordinated at different developmental stages and constraints (i.e. larval-homeostasis, pupal endosymbiont-reinfection, adult-contrasted endosymbiont dynamics), we have conducted a host-symbiont metatranscriptomic approach (dual RNA-seq), and obtained high-throughput data simultaneously from both partners throughout 12 key stages of the weevil development. To pinpoint endosymbiont-dependent changes in gene expression during young adult contrasted bacterial dynamics, we performed an additional comparative transcriptomic analysis between symbiotic and aposymbiotic insects. We show that larval stages are not as transcriptionally stable as anticipated, marked by an enrichment in genes involved in cell growth from both host and bacteria. Following bacteriocyte migration, and while insects undergo metamorphosis, *S. pierantonius* activates secretion system and virulence genes, concomitantly with stem cell infection, bacteriocyte differentiation and antimicrobial peptide (AMP) encoding genes induction by the host. Metabolic and regulatory genes involved in this process are coordinated in both insect and bacteria. We identified several signaling pathways, such as Hippo, Wnt and Notch along with iron metabolism as symbiotic-specific features and likely essential for bacterial growth. Once the bacterial benefit is over, insects activate recycling enzymes along with genes involved in apoptosis and autophagy. In response, bacteria activate stress-related-genes including all cytoplasmic chaperones.

Overall, our findings show that an interkingdom dialogue and coordinated regulation of gene expression play a pivotal role in the morphological reorganization of the bacteriome and the bacterial dynamics from larval stages to adulthood, highlighting adaptive features and tightly regulated biological functions to support both bacterial growth and recycling in emerging adults.

## Material and methods

### Animal rearing and sampling

In this study we used the *S. oryzae* strain Bouriz, which is *Wolbachia*-free and bears *S. pierantonius* exclusively (Figure 1A). Insects are reared on wheat grains at 27.5 °C and at 70 % relative humidity. Aposymbiotic animals were obtained by heat treatment and have been maintained in laboratory conditions similarly to the symbiotic line[39]. For the dual RNA-seq experiment, we sampled 12 time points with three biological replicates for symbiotic insects during the insect development, namely: 3rd and 4th instar larvae (L3 and L4 respectively), early pupae (EP), pupae (P), along with eight time points into young adulthood starting from Day 1 to Day 20: D1, D2, D3, D5, D7, D9, D12, D20 (Figure 1B). Five time points were sampled for the regular RNA-seq in both symbiotic and aposymbiotic animals: D1, D3, D5, D7, and D9 (Figure 1C), with triplicates except for symbiotic D5 samples, which had four biological replicates. Bacteriomes were dissected in diethylpyrocarbonate-treated Buffer A (25 mM KCl, 10 mM MgCl_2_, 250 mM sucrose, 35 mM Tris/HCl, pH = 7.5). For each replicate, bacteriomes (from 40 L3 and 30 L4) or midguts (from the remaining time points, ∼15 insects for dual RNA-seq libraries, and 8-10 insects for each RNA-seq sample) were pooled, and stored at −80 °C before RNA extraction.

**Figure 1.**
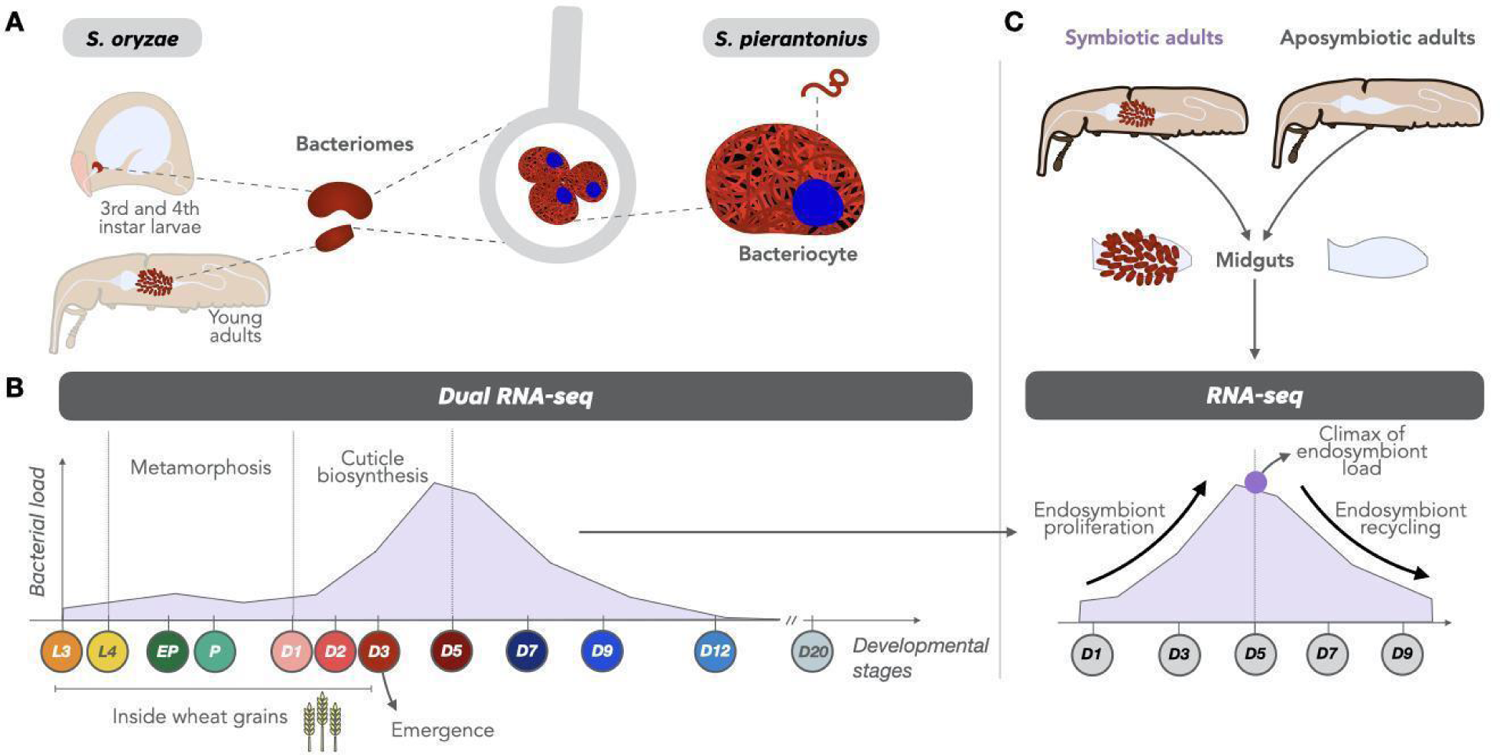
Experimental design for deciphering the molecular dialogue between *Sitophilus* oryzae and its endosymbiont Sodalis pierantonius. (A) Schematic illustration of the bacteriome location in larvae and adult weevils. (B) Key developmental life stages sampled in this study for the dual RNA-seq and schematic of gut bacterial load from previous studies.[40,41] (C) Time points sampled for the RNA-seq comparison of symbiotic and aposymbiotic weevils. Arrows depict the increase in bacterial load during the first days of adulthood, followed by the drastic recycling of endosymbionts. L: larvae, EP: early pupae, P: pupae, DX, estimation of the adult age in days.

### RNA extraction, library preparation and sequencing for dual RNA-sequencing

Total RNA was extracted with TRIzol™ Reagent (Invitrogen ref.: 15596026) following the manufacturer’s instructions. Nucleic acids were then purified using the NucleoSpin RNA Clean up kit (Macherey Nagel ref.: 740948). Genomic DNA was removed from the samples with the DNA-free DNAse kit (Ambion ref.: AM1906). Total RNA concentration and quality were checked using the Qubit Fluorometer (ThermoFisher Scientific) and Tapestation 2200 (Agilent Biotechnologies). Ribo-depletion was performed by using a total of 170 rRNA depletion probes designed using the Ovation Universal RNA-seq System (NuGEN) following the manufacturer’s instructions (kit reference: 0343-32) in order to target all prokaryotic and eukaryotic rRNA sequences from both *S. pierantonius* (5S, 16s and 23S) and *S. oryzae* (5.8S, 18S, 28S and 12S and 16S mitochondrial rRNAs). Dual RNA-seq libraries were then generated with 100 ng of total RNA and sequenced on a NextSeq 500 instrument (Illumina) at the sequencing platform of the Institut de Génomique Fonctionnelle de Lyon, Ecole Normale Supérieure de Lyon, France. Supplementary Figure S1A explains the schematic pipeline for the dual RNA-seq.

### RNA extraction for host RNA-sequencing

Total RNA was extracted using the Qiagen Allprep DNA/RNA micro kit (Qiagen ref.: 80284) following the manufacturer’s protocol. RNA quality assessment and library preparation were performed at the GenomEast platform at the Institute of Genetics and Molecular and Cellular Biology using Illumina Stranded mRNA Prep Ligation (Reference Guide - PN 1000000124518). RNA-Seq libraries were generated according to manufacturer’s instructions from 200 ng of total RNA using the Illumina Stranded mRNA Prep, Ligation kit and IDT for Illumina RNA UD Indexes Ligation. Libraries were sequenced on an Illumina HiSeq 4000 sequencer as paired-end 100 base reads.

### Preprocessing, mapping of reads and differential expression analysis

Supplementary Figure S1B depicts the processing and bioinformatic analyses. The raw reads from both sequencing methods (dual RNA-seq and RNA-seq) were processed using CUTADAPT v1.18[42] to remove adapters, and filter out reads shorter than 50 bp and reads that had mean quality of 30 or less. The clean reads were mapped against the *S. oryzae* genome (GCF_002938485.1) with STAR v2.7.3a[43] and S*. pierantonius* (CP006568.1) with BOWTIE v2.3.5[44] when suitable. Supplementary Table S1 presents all information regarding the number of raw reads, trimming and mapping to each of the genomes (when suitable) for both sequencing methods. Shared reads from dual RNA-seq between the two genomes were filtered out with the aid of SAMTOOLS[45] and PICARD (available from https://broadinstitute.github.io/picard/). The high proportion of multi-mapped reads from both genomes is explained by the high abundance of repeats in both *S. oryzae* [46] and *S. pierantonius* [19] (74% of repeats in *S. oryzae* and 19% of insertion sequences in *S. pierantonius*). Gene counts were obtained for uniquely mapped reads with FEATURECOUNTS v2.01 method from the SUBREAD package[47] (Supplementary Tables S2 and S3 for host and endosymbiont dual RNA-seq results respectively). The SARTOOLS package[48] was used for quality assessment of samples and pairwise differential expression analyses using DESEQ2 v1.34.0[49]. After testing, the p-values were adjusted with the Benjamini-Hochberg[50] correction for multi-testing. Genes were considered differentially expressed when adjusted p-values < 0.05 (results for differential expression of the dual RNA-seq data are provided in Supplementary Tables S4 and S5). Each time point was tested against all time points to construct an overall gene expression graph, however, we looked closely at the results between two successive time points (*t* versus its previous time point *t - 1*). Since we had a high number of conditions and pairwise comparisons tested within the dual RNA-seq dataset, we also performed a likelihood ratio test (LRT) to create a reduced model and detect genes showing the most significant variation across all time points (Supplementary Table S6 and S7). For the regular host RNA-seq, counts are provided in Supplementary Table S8. A comparison with the previous time point, as well as a linear model, were performed with DESEQ2 using the following design: ∼time+symbiont+interaction (Supplementary Table S9). Variability between samples intra-condition was verified with dispersion metrics from DESEQ2. Supplementary Text S1 provides information on differential expression results for both datasets.

### Clustering and profiling analyses

The standardized expression (normalized to mean zero and standard deviation one) of differentially expressed genes (DEGs, adjusted p-value < 0.05) of both insect and bacteria were loaded into MATLAB v9.12.0 R2022a and separately clustered using a temporal clustering by affinity propagation with the method TCAP v.2[51]. The results were then loaded and analyzed in R v4.1.2[52] and clusters with at least 0.8 of mean similarity which presented similar profiles but differed mainly by the magnitudes of expression were merged into superclusters using Euclidean distance with the PHEATMAP package v.1.0.12[53]. Clusters with less than 10 genes were discarded and visualization of profiles was made with GGPLOT2 package v3.3.6[54]. Supplementary Tables S10 and S11 present the clustering results for the host and endosymbiont respectively from the dual transcriptomics dataset and Supplementary Table S12 presents the clustering results for the host RNA-seq data between symbiotic and aposymbiotic adults.

### Functional enrichment analysis

Functional analysis took into account as input the lists of DEGs and superclusters. We performed a global functional enrichment analysis using Cluster Profiler[55] with multi-testing correction[50] in R to find out which functions were over-represented in each comparison. Gene Ontology[56] (GO) terms and KEGG[57] pathways with adjusted p-values < 0.05 were defined as significantly enriched. We also visualized biological pathways that were highly impacted in both insect and bacteria, by projecting the transcriptomic data onto the online platform Search and Color from KEGG. When needed, GO terms were reduced to representative non-redundant terms with the use of the REVIGO tool[58]. Results for the dual RNA-seq dataset are provided in Supplementary Tables S13 and S14 for both host and endosymbiont. Results for the RNA-seq dataset are provided in Supplementary Table S15.

## Results and Discussion

One of the typical features of weevil endosymbiosis (Figure 1), is the contrasted endosymbiont dynamics in young adults. Following adult metamorphosis and gut stem cell infection, the bacterial population grows exponentially for a few days, concomitantly with the biosynthesis of the insect’s cuticle, and soon after the bacteria are actively recycled by the host. At 10-12 days of adulthood, laboratory-reared insects are almost entirely devoid of midgut bacteria[40]. Thus, at least three distinct developmental constraints can be distinguished after metamorphosis: initial bacterial growth concomitant to bacterial benefit, climax of bacterial load, and subsequent symbiont elimination once bacterial burden overcomes its benefits.

To identify host and bacterial genes and pathways involved in bacterial growth regulation, we have established a host-symbiont transcriptomic approach (dual RNA-seq), resulting in high-throughput data simultaneously from both partners. We have analyzed insect and bacterial gene expression at 12 key stages of the weevil development, from 3rd instar larvae to 20-day-old adults (Figure 1B). To pinpoint endosymbiont-dependent gene expression, we performed an additional transcriptomic comparison between symbiotic and artificially obtained aposymbiotic insects (Figure 1C).

The number of differentially expressed genes (DEGs) varied drastically in both weevils and endosymbionts (Supplementary Text S1, Supplementary Tables S4 and S5). For insect genes, DEGs ranged from 38 (between D12 vs D9) to 6229 (between D1 vs P), as presented in Table 1. Due to extreme tissue rearrangements, a major shift in expression observed during metamorphosis had already been reported between L4 vs EP developmental stages[36]. Here, we monitored an even higher number of DEGs than previously, especially regarding the endosymbiont, thanks to the deeper sequencing obtained for this study. After the bacterial elimination period (D12 and D20), the expression of insect genes is stable, with nearly no DEGs between D12 vs D9, nor D20 vs D12. As for the endosymbiont, we detected an overall correlation between bacterial load and bacterial gene expression (Supplementary Figure S2) and a maximum of 468 genes were differentially expressed (between D1 vs P). Even between L3 vs L4 larval stages, which were considered homeostatic due to the coordinated growth of the bacterial population and the bacteriome tissue, a significant number of DEGs from both partners was observed, attesting that *S. pierantonius* gene expression is differentially regulated and responds to different developmental conditions.

**Table 1.**
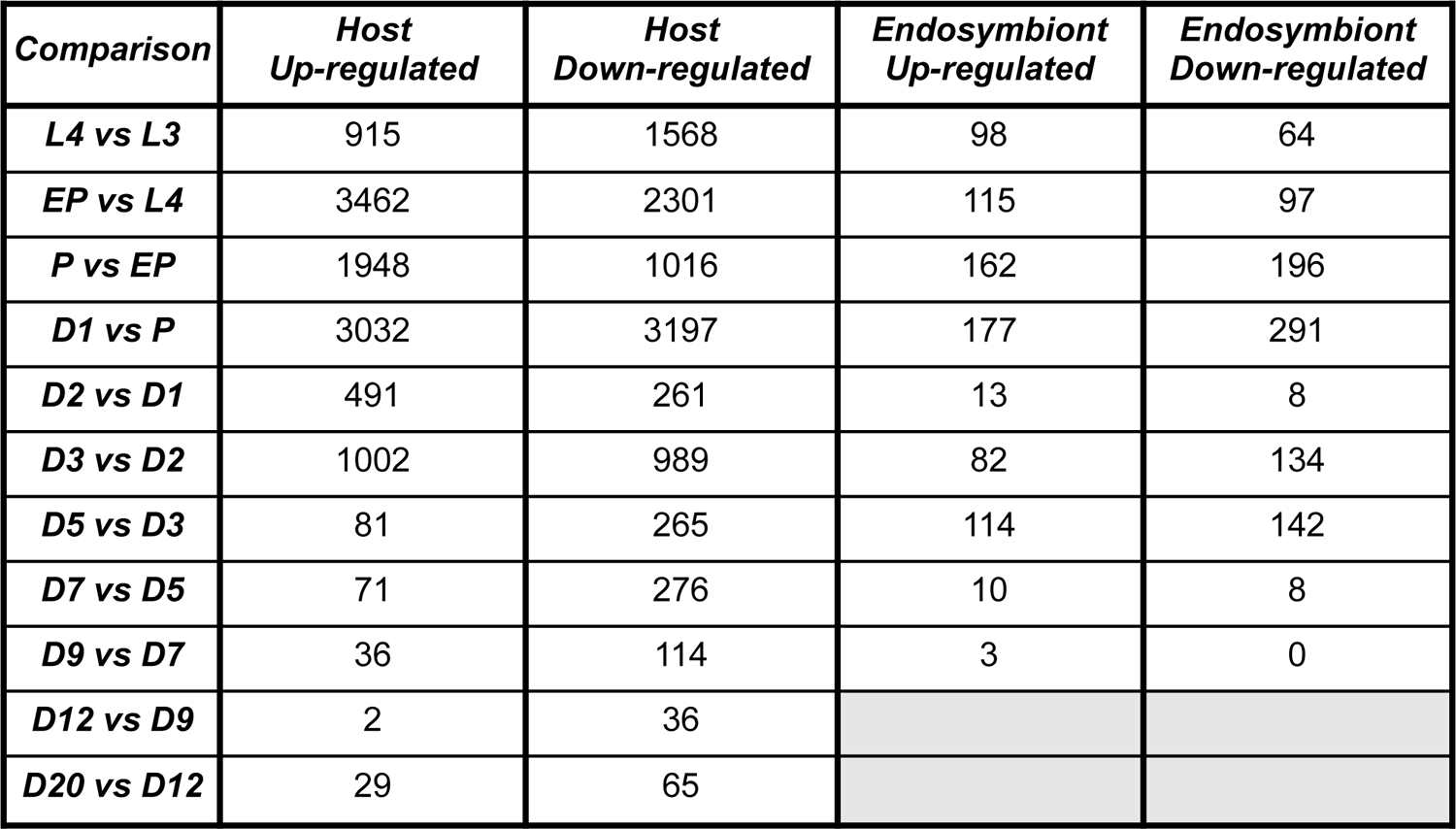
Number of DEGs detected for *S. oryzae* and *S. pierantonius* throughout 12 time points during the insect’s life cycle. Genes were considered as differentially expressed when the adjusted P-value was lower than 0.05 and the absolute Log2 of fold change was greater than 0.5.

Based on the centered normalized expression of DEGs, we conducted a clustering analysis of genes presenting similar profiles with TCAP2[51]. Finally, clusters of genes having similar expression but different magnitudes were further merged into superclusters (Supplementary Tables S10 and S11). The overall results of selected superclusters throughout the developmental stages from both host and endosymbiont are depicted in Figure 2 (Supplementary Figures S5 and S6 provide complete profiles of all superclusters). Superclusters composed of genes from the insect will be denoted as **SO** followed by the supercluster number; whereas superclusters composed of genes from the endosymbiont will be denoted with an **SP** followed by the number of the supercluster. Functional enrichment of the final superclusters (Supplementary Tables S13 and S14) unraveled several biological functions during most of the stages of the developmental process and will be discussed in the following subsections. The main enriched functions associated with changes in insect or bacterial gene expression are depicted in Figure 2. Gene names or abbreviations mentioned throughout this manuscript are provided with the respective gene IDs in Supplementary Table S16.

**Figure 2.**
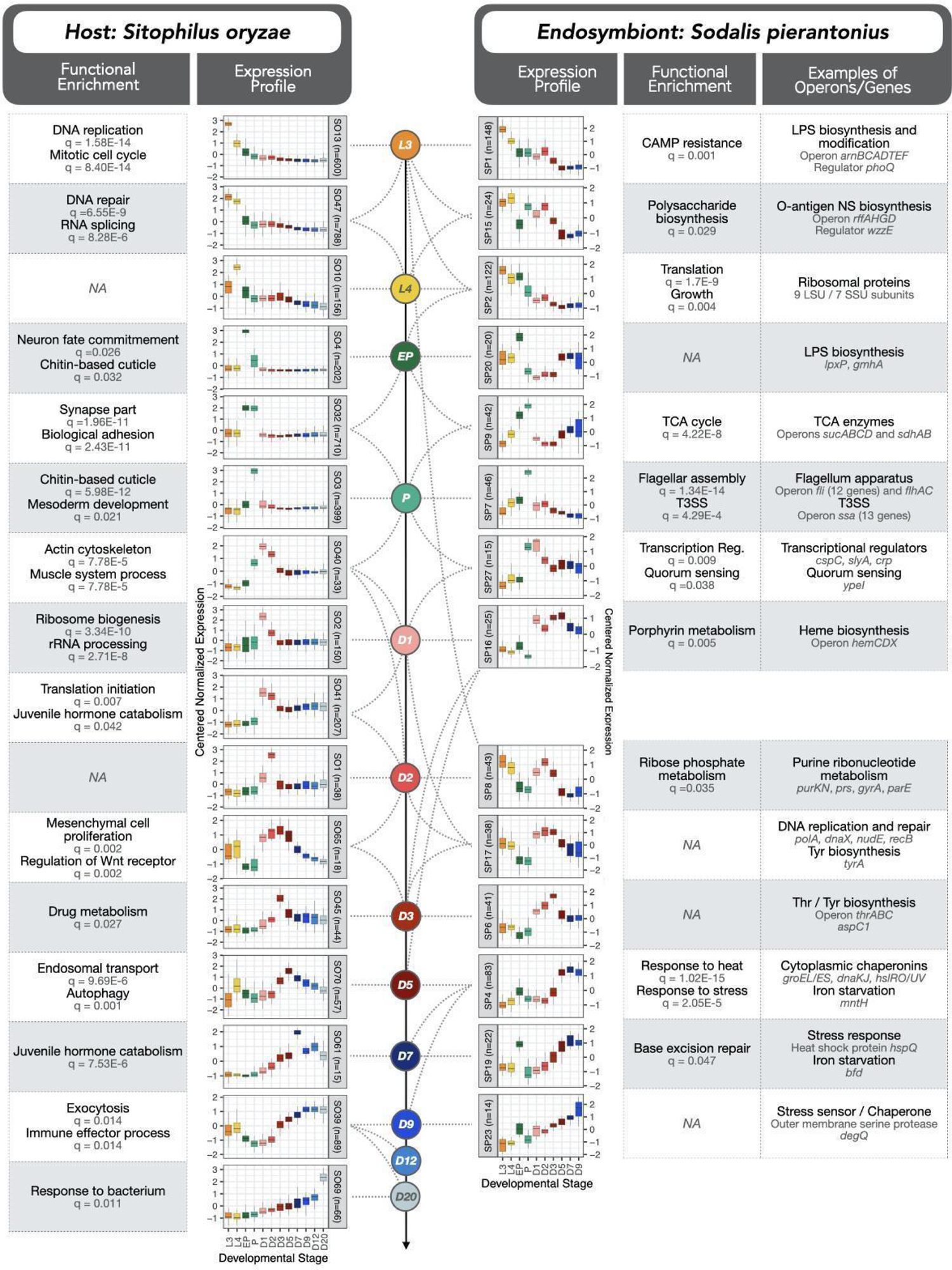
Dual RNA-seq expression profiles of both host and endosymbiont during the 12 stages of development studied. Profiles highly induced in each of the stages (or in more than one stage at times) are indicated with dashed lines. Host profiles are indicated with SO (for *Sitophilus oryzae*) followed by the profile number whereas endosymbiont profiles are indicated with SP (for Sodalis pierantonius) followed by the profile number. Each profile was built based on the expression of different numbers of genes (indicated by n). Color for each developmental stage follows the same color coding as Figure 1. Functional enrichment of each profile based on GO and KEGG is indicated on the left side for host genes and on the right for endosymbiont genes. We also provide, on the far right, examples of endosymbiont genes belonging to each profile. The complete set of profiles built is provided in Supplementary Figures S5 and S6.

### Larval stages show active cell growth in both host and endosymbiont

When compared to pupal and adult stages, genes highly expressed in larval bacteriomes show a significant functional enrichment related to mitosis, nuclear division, mitochondrion, ribosome, RNA splicing, nucleolus, DNA repair, and protein refolding, suggesting a high proliferation of cells, especially at the L3 stage (superclusters SO13, SO47 and SO10 from Figure 2; SO28 from Supplementary Figure S3). These results support the idea that the larval bacteriome is in a growing phase, presumably storing up energy and metabolites before insects stop feeding and start metamorphosis. The gene *tor,* a serine/threonine protein kinase that regulates and promotes cell and tissue growth in *Drosophila*[59], is highly induced at L4 stage (supercluster SO10), along with bacterial genes involved in ribosome synthesis and translation, gene expression, growth, cell wall, and lipopolysaccharide (LPS) biosynthesis (superclusters SP1, SP2 and SP15), concomitantly with a slight increase in bacterial load from L3 to L4. The most significant gene from the whole bacterial dataset is an LPS biosynthesis gene, namely *murB* (p-adj = 1.09E-277, Supplementary Table S7), whose expression is high at L3, and drops more than 40-fold by D9. A similar pattern of active cell growth coordination has been reported during the host proliferative state in aphids, characterized by larger and more numerous bacteriocytes[28].

The only AMP highly expressed in larval bacteriomes, and whose expression parallels the profile of bacterial load in adult stages, is a partial Diptericin-like protein coding gene (*dpt-like partial*, belonging to cluster SO47). This gene, unlike the canonical *diptericins*, lacks the 5’ end that encodes a signal peptide for export[46]. This AMP is predicted to be intracellular and likely resides within bacteriocytes, similarly to the Coleoptericin A (ColA), an AMP that was already shown in weevils to continuously target the endosymbionts and inhibits their cell division[11,60]. A complete bacterial operon (*arn*) acting on LPS modification and involved in AMP resistance in *Dickeya dadantii*[61], was induced at larval stages (supercluster SP1) (See Supplementary Text S2A for further information, refs[11,60–63]).

Six bacterial genes from the ATP synthase operon (*atpEAGDBH*) also belong to the superclusters SP1, SP2, SP8, and SP15, showing high expression in larval stages. Previous biochemical studies have reported that weevil mitochondrial-specific enzymatic activities are higher in symbiotic larvae than in aposymbiotic larvae[64,65]. Here, we provide further evidence that the increase in host mitochondrial activity is part of a coordinated regulation between bacteria and host to generate energy necessary for an active metabolism within the bacteriocytes and endosymbiont cells.

Finally, it has been recurrently suggested that the larval bacteriome is at a homeostatic state between the host and endosymbiont, since bacterial growth is not significantly important, bacteriocytes do not undergo cellular rearrangements, and the host immune response is down-regulated, with the notable exception of ColA[13,36]. Here, we show that both the host and endosymbiont actively ‘prepare’ for the physiological and energetic needs necessary for insect metamorphosis, along with bacteriocyte migration and differentiation of new bacteriomes during the pupal stages. In sum, while the larval bacteriome exhibits a homeostatic status regarding bacterial population size and bacteriocyte cellular processes, this work reveals that host and bacterial coordinated gene expression precedes energy production and component synthesis necessary for host metamorphosis execution and completion.

### Metamorphosis and AMP expression depict an arms race between symbiotic partners

During metamorphosis, the single larval bacteriome disassembles, bacteriocytes migrate along the midgut, and bacteria infect new stem cells at the apexes of midgut caeca, which differentiate further into novel bacteriocytes that group into several bacteriomes around the midgut of the young adult[36]. As expected, and per our previous observations, insect genes highly expressed in pupal stages were overall enriched in cell adhesion, mesodermal cell commitment, chitin based-cuticle and synapse (superclusters SO3, SO4, and SO32 in Figure 2; SO9 and SO11 from Supplementary Figure S3). Additional insights of the insect gene, *tolA*, that could directly impact bacterial gene expression during metamorphosis are provided in Supplementary Text S3A (ref[66]).

Concomitant to the insect’s metamorphosis, the bacteria showed an increased expression in TCA cycle, and several virulence-related genes (superclusters SP7 from Figure 2, SP10 from Supplementary Figure S4), in agreement with our previous work[36]. Virulence genes included the partially degraded operons of the flagellum apparatus (operons *fli* and *flh*), and type III secretion system (T3SS, operon *ssa* and *sseE*). No apparent effector for the T3SS was ever reported in *S. pierantonius*. Here, we have detected a pseudogene highly expressed in P stage exclusively, homolog to the *Salmonella typhimurium* deubiquitinase SseL, a T3SS known effector[67] (Supplementary Figure S5). Despite a premature stop codon, the *S. pierantonius sseL* gene presents an intact peptidase domain. SseL inhibits autophagic clearance of cytoplasmic aggregates that form during *Salmonella* infection in epithelial cells, favoring intracellular bacterial replication[68]. The upregulation of *sseL* during weevil’s metamorphosis suggests an essential role in the infective behavior of *S. pierantonius*.

We have also detected a spike in 10 insect AMP genes during metamorphosis, namely *colA*, *coleoptericin-B*, *acanthoscurrin-1-like*, *cathelicidin-like antimicrobial protein*, *cecropin*, *diptericin-2*, *diptericin-3*, *diptericin-4*, *gly-rich AMP-like*, and *holotricin-3-like*, (Supplementary Figure S6). At first, it is appealing to hypothesize that this increase in AMP expression concomitant to the process of bacterial infection of stem cells could be a response of the host to the virulent bacteria exiting the bacteriocytes. However, recent work from our group has looked further into this peak of expression of AMPs using RT-qPCR during the metamorphosis and showed that the expression of AMPs is independent of endosymbiont presence[69]. Indeed, even though AMP genes can be induced upon infection, there is growing evidence suggesting they are activated at specific developmental stages regardless of the presence of a bacterium[70,71].

In conclusion, contrary to *colA*, nine AMPs were shown to be upregulated precisely during metamorphosis endosymbiotic-independently, suggesting that novel immune regulators might be involved in a developmental-specific manner.

### Early stages of adulthood depict active tissue and bacterial growth and coordinated metabolism

As we previously mentioned, there are three important features of bacterial dynamics tightly regulated during the young adult developmental stages: endosymbiont proliferation, climax in bacterial load and subsequent bacterial recycling. We recently showed that endosymbiont proliferation relies on metabolic regulation and the availability of carbohydrates from diet intake, and this proliferation seems to escape host control[41]. In contrast, both the bacterial load climax and subsequent recycling were described to be genetically controlled by the host[40,72], even though the specific genes orchestrating these features remain unknown. Thus, our main focus in the following sections was to identify genes and pathways involved in bacterial dynamics, along with the coordinated bacterial responses to host changes.

Host genes enriched at the initial stages of adulthood (D1 and D2), before insect emergence from the grain (superclusters SO1, SO2, SO40, SO41, from Figure 2; SO16, SO54, and SO55 from Supplementary Figure S3), had a significant functional enrichment related to ribosome biogenesis, rRNA processing, actin cytoskeleton, muscle system process, suggesting tissue development along with cellulose degradation in both insect and bacteria.

Bacterial gene enrichment analysis (superclusters SP8, SP16, SP17) showed a host-endosymbiont coordination in the biosynthesis and metabolism of other precursors necessary for growth (Supplementary Text S4, refs[19,41,46,73–75]). This includes the activation of DNA replication and repair along with the biosynthesis of heme and metabolism of ribonucleotides from D1 to D3. Particularly at D1, we noticed an increased expression of bacterial transcription factors (supercluster SP27), and possibly a general transcriptional activation, suggesting that even though the insect is not feeding until shortly before emergence from the grain, the bacteria seem not affected by this lack of resources, relying on stored metabolic components during the previous larval stages. From D3 to D5, the bacteria proliferate inside the growing bacteriocytes and actively produce essential metabolites necessary for the host[46]. Metabolism is usually regulated straightforwardly, in which the availability of a substrate or the lack of a product in a certain pathway induces the expression of key genes in such a pathway. This regulation becomes more complex when dealing with two distinct organisms, coming from different kingdoms of life, and two examples of these exchanges are the essential amino acids threonine and tyrosine, provided to the host by the endosymbiont[40,46,76]. We show that the operon *thrABC* (threonine synthase) and the genes *aspC1* and *tyrA* (tyrosine biosynthesis) are induced after metamorphosis with a peak at D2/D3 (superclusters SP6 and SP17). This suggests that the provision of such essential amino acids along with other cofactors from the endosymbiont to the host initiates before the insect leaves the grain and likely helps the insect before cuticle synthesis and mouthparts acquisition during the fasting period. The tyrosine amino acid is particularly critical since it is one of the main precursors of the dihydroxyphenylalanine (DOPA), an essential component of the insect’s cuticle synthesi[40], which is fully completed around D5 in weevils. A supercluster from the host with a higher expression from D1 to D5 (SO65), had enriched terms related to mesenchymal cell proliferation and stem cell maintenance, suggesting gut and bacteriome tissue growth until the cuticle is completed. Interestingly, a final up-regulation of genes belonging to the juvenile hormone catabolism was detected around D7 (an example is the carboxylesterase *vest-6*, supercluster SO61), likely depicting the insect’s maturation after the completion of the cuticle.

### Transcriptomic profiling of symbiotic and aposymbiotic adults revealed an endosymbiont-driven regulation

To uncover host genes affected by the endosymbiont presence, we performed a regular eukaryotic RNA-seq between gut tissues dissected from symbiotic and aposymbiotic insects at five key stages of young adulthood (D1, D3, D5, D7, and D9, as presented in Figure 1B). We performed a differential expression analysis and detected that the presence of the endosymbiont was the major effect controlling either the induction or the depletion of gene expression throughout the dataset (Figure 3). The AMP gene *dpt-like*, for instance, which is highly prevalent in bacteriome tissue from larval stages (Figure 2), is highly induced at D1 and D3 only in symbiotic insects. Another interesting example of a gene following the bacterial climax codes for an Anillin-like protein (Figure 3), a scaffolding protein known to regulate integrity of intercellular junctions[77–79], and which nonetheless could play a role in the overall stability of the bacteriomes.

**Figure 3.**
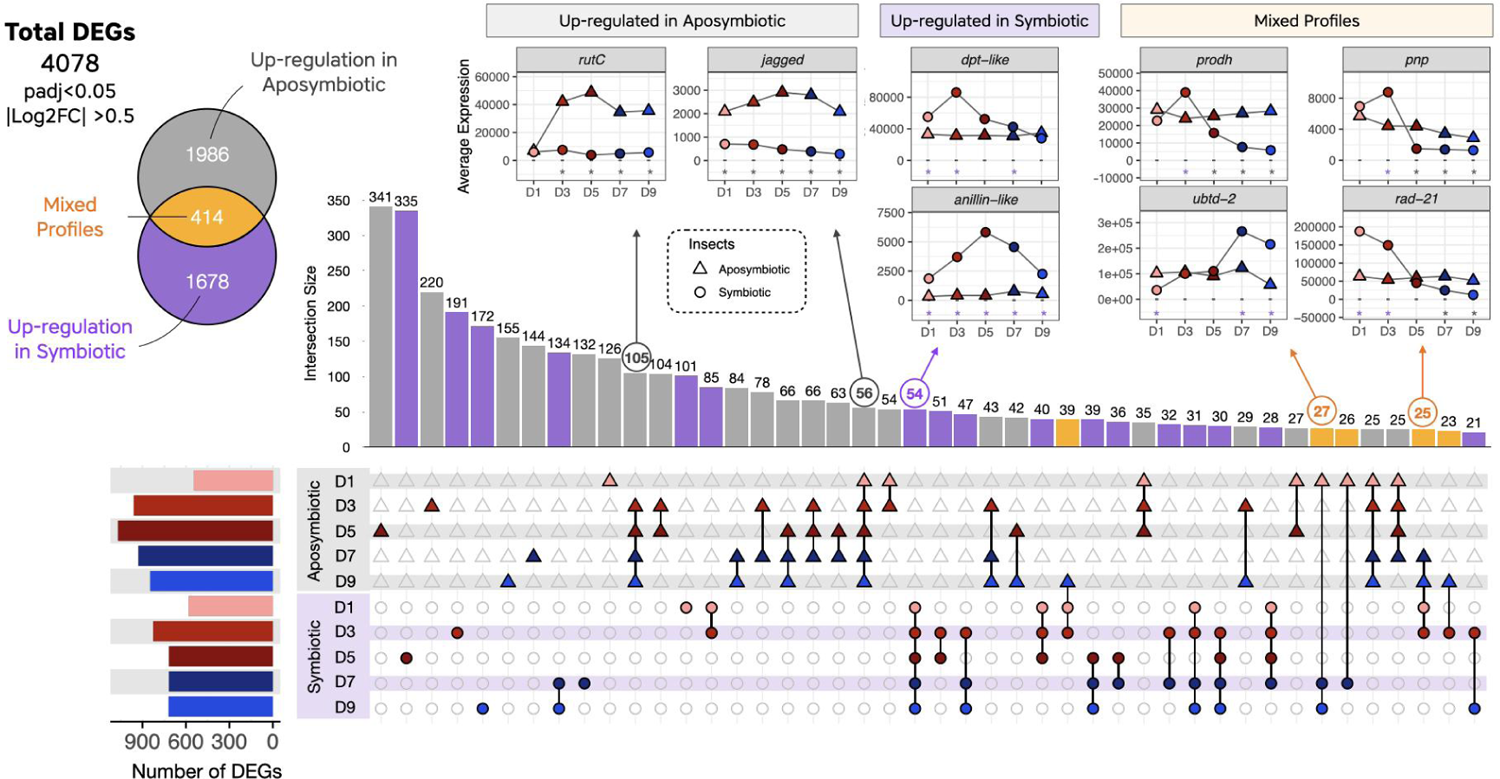
UpSet plot with overview of transcriptomic profiling between symbiotic and aposymbiotic strains. In total, we detected 4078 genes differentially expressed in at least one of the five developmental stages between symbiotic and aposymbiotic insects. Around half of these genes were differentially expressed stage-specifically (1917 in total, either as single dots or single triangles respectively for symbiotic or aposymbiotic strains), 1747 were either up-regulated in more than one condition from symbiotic (purple) or aposymbiotic (gray) insects, while 414 genes presented mixed profiles (in yellow). To better illustrate DEGs common to more than one stage, we present on the top of the UpSet plot some examples of genes belonging to different UpSet groups. Stars indicate in which stages we detected a significant up-regulation (purple) or down-regulation (gray) in symbiotic insects when compared to aposymbiotic within the RNA-seq dataset.

Similarly to the dual RNA-seq analysis, we combined clusters from symbiotic and aposymbiotic insects into Superclusters (Figure 4, and Supplementary Figures S7 and S8), by taking into account their expression profiles and the stages in which we detected a differential expression between symbiotic and aposymbiotic weevils (Supplementary Table S10). Functional enrichment was performed (Supplementary Table S15) and superclusters induced in symbiotic adults will be denoted with an **S**, whereas superclusters induced in aposymbiotic insects with an **A**. Insights on genes and enriched functions from aposymbiotic insects are provided in Supplementary Text S3B (refs [80–82]).

**Figure 4.**
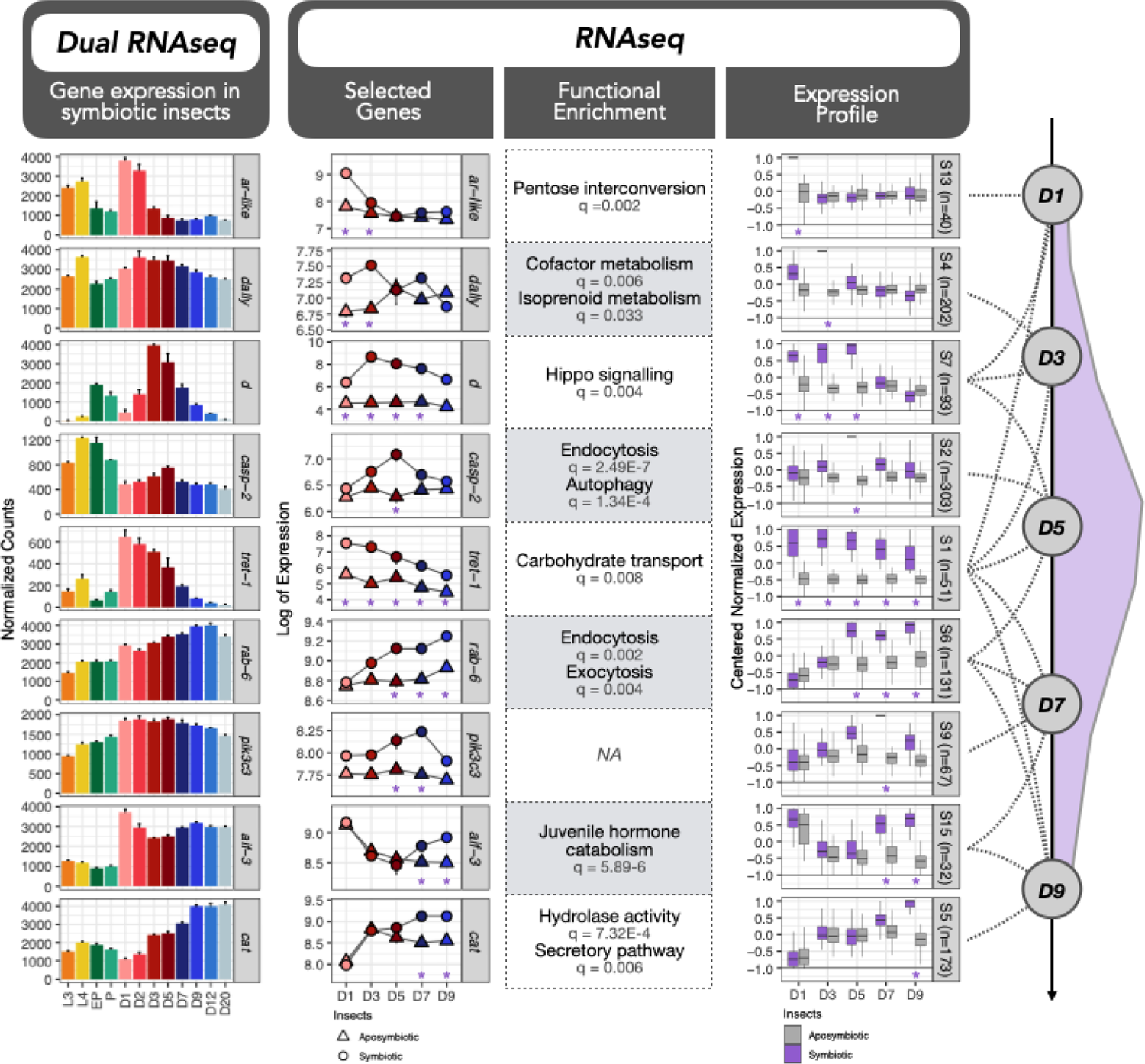
Clustering of genes differentially upregulated in symbiotic insects during the young adult bacterial dynamic. Gene expression of selected genes are given for the dual RNA-seq (first panel) and host RNA-seq (selected genes, RNA-seq panel) as examples. Color for the dual RNA-seq dataset follows the same color coding as Figure 1. Expression profiles of clusters of genes highly prevalent in one or in more than one stage in symbiotic insects are indicated with dashed lines and are indicated with S followed by the profile number. Each profile was built based on the expression of different numbers of genes (indicated by *n*). Purple color indicates symbiotic insects whereas gray indicates aposymbiotic insects. Purple stars indicate in which stages we usually detected significant up-regulation in symbiotic insects when compared to aposymbiotic insects within the RNA-seq dataset. Functional enrichment of symbiotic superclusters provide functions specifically regulated in symbiotic weevils. On the far right, the bacterial load during the young adult development is shown in purple.

In agreement with the findings that endosymbiont proliferation relies on the availability of carbohydrates from food intake[41], genes upregulated in all symbiotic stages were enriched in transport of carbohydrate functions (Supercluster S1; Figure 4). Interestingly, pentose interconversion and cofactor metabolism genes were induced specifically in symbiotic insects at D1 and D3 (Superclusters S13 and S4), which presumably helps to provide essential substrates for the imminent bacterial growth[46]. It is noteworthy that at D1 the insect is still fasting, suggesting that this provision of sugars and cofactors to the endosymbiont comes from larval reserves. When searching for genes induced in specific stages in symbiotic strains, we detected similar functional enrichment (Figure 4) to the dual RNA-seq dataset presented in Figure 2 (such as exocytosis, endocytosis, autophagy, juvenile hormone catabolism, for further information see Supplementary Text S3C, refs[83,84]), along with others specific to this dataset (such as Hippo signaling).

### Host signaling pathways governing bacterial dynamics

Previous studies on endosymbiont maintenance and control indicate that chronic infection of intracellular bacteria in insects interferes with a broad range of cellular functions[85], including oxidative stress[86], signaling cascades[28], immunity[11,87], apoptosis and autophagy[40,88].

In this view, we compiled all annotated genes from regulatory pathways[46] differentially expressed either throughout the development of the cereal weevil or differentially expressed between symbiotic and aposymbiotic insects (Supplementary Table S17) in search for symbiotic-specific regulatory mechanisms. The strongest enrichment within signaling pathways was identified within the symbiotic/aposymbiotic RNA-seq dataset, regarding genes from the Hippo signaling cascade pathway at D1, D3 and D5 (Supercluster S7, Figure 4). We found 43 genes (out of 49 homolog genes from this pathway in *Drosophila*), differentially expressed between symbiotic and aposymbiotic weevils, most of which were upregulated in symbiotic insects.

From both transcriptomic datasets (Figure 5), we show that two repressors of Warts activity (*d* and *ds*) followed a symbiotic and time-dependent peak of expression at D3, and four activators of Hippo were induced in symbiotic insects either in a more stable manner up until D5 (*kibra*, *ex*) or with peak of expression precisely at D5 (*mer*, *sav*), suggesting a transcriptional activation of this pathway until D5, when the bacterial load climax is reached. In aphids, the Hippo pathway is up-regulated in host proliferative states also in response to essential amino acid limitation[28,89]. Given that the regulation of this pathway relies heavily on post-transcriptional regulation (mostly phosphorylation and proteolysis) and that symbiotic and aposymbiotic weevils share the same genetic background as symbiotic ones, it is surprising to detect this activation of the Hippo signaling at the transcriptional level only in symbiotic weevils.

**Figure 5.**
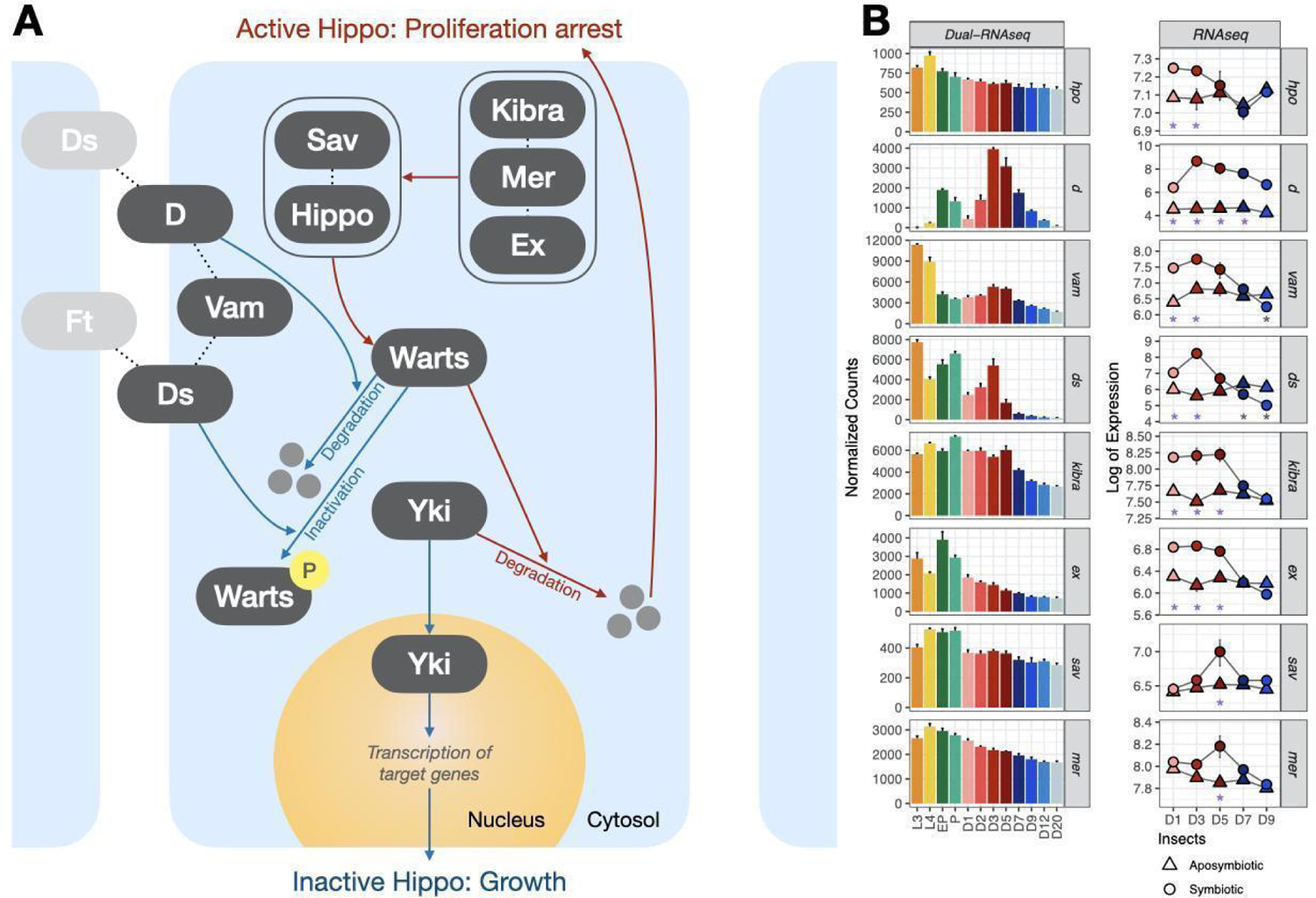
Hippo signaling in *S. oryzae*. A. Simplified diagram of the Hippo signaling pathway proposed in the cereal weevil B. Genes involved in Hippo signaling that were detected as differentially expressed throughout the life cycle of the cereal weevil (dual RNA-seq) and/or differentially expressed between symbiotic and aposymbiotic insects (RNA-seq). When the Hippo signaling pathway is inactivated, the transcriptional co-activator Yorkie (Yki) enters the nucleus and induces transcription of target genes, leading to growth. When the Hippo signaling is activated, Warts kinase phosphorylates Yorkie, inducing its ubiquitination and degradation, therefore leading to a proliferation arrest. Hippo, Salvador (Sav), Kibra, Merlin (Mer) and Expanded (Ex) are examples of activators of the Hippo pathway, while Dachshous (Ds) and Dachs (D) are examples of repressors of Warts activity. Stars indicate in which stages we detected a significant up-regulation (purple) or down-regulation (gray) in symbiotic insects when compared to aposymbiotic within the RNA-seq dataset.

Moreover, the protein Ds is also involved in the non-canonical Wnt signaling pathway, which has already been hypothesized to play a role in bacteriocyte differentiation and bacteriome formation in weevils[90]. Interestingly, a significant enrichment of “Regulation of Wnt receptor” was detected at the initial stages of adulthood (Supercluster SO65, Figure 2). Besides *ds*, we found ten genes belonging to the Wnt signaling pathway that presented differential expression profiles, five of which (*mwh*, *daam*, *nmo* and two loci for *dally*) are induced in symbiotic weevils at D1 and D3 (Supercluster S4). The Wnt signaling pathway has been proposed to participate in the communication between bacteriocytes, body and bacterial cells in aphids[91], and both Wnt and Hippo have been implicated in the growth regulation of disproportionally growing organs in the hemimetabolous cricket *Gryllus bimaculatus*[92]. Genes belonging to the Notch signaling pathway were also detected as differentially expressed between symbiotic and aposymbiotic insects (See Supplementary Text S3D, refs[93–96]).

In sum, these results highlight specific regulatory factors from Hippo and Wnt pathways that might be central in the regulation of bacterial load in the cereal weevil. Nevertheless, further studies are needed to investigate whether these signaling pathways regulate the increase in bacteriome size during the first days of adulthood, the bacterial load climax achieved around D5, or pre-signaling the following period of bacterial recycling and cell death.

### Nutritional immunity seems to be central in the control of the endosymbiont load

Nutritional immunity, or the control over transition metals, is one of the first lines of defense against bacterial infections[97]. Iron, for example, is essential for all living organisms to transport oxygen and produce energy. Nevertheless, since iron ions are capable of catalyzing destructive oxidative reactions, organisms have evolved various strategies for controlling and sequestering it[98]. Thus, bacteria have evolved parallel pathways to recover iron from different sources when starved for iron[99]. We have identified genes related to the sequestration of transition metal ions with profiles that showed an inverted (Supercluster SO41, Figure 2), or direct correlation with the bacterial load (Supercluster SO45 and SO70, Figure 2).

In this respect, we have evaluated the expression profile of several genes involved in the metabolism, uptake and storage of metals from both insect and bacteria (Figure 6). Two proteins that can sequester iron and keep it from a destructive association with oxygen are transferrins (Tsf1) and ferritins (Ft)[98]. Within the dual RNA-seq data, these proteins showed divergent transcriptomic profiles: while transferrin (*tsf-1*) had a profile similar to the inverted bacterial load (supercluster SO41), ferritin heavy chain-like (*fth-1*) expression peaked at D3 (supercluster SO45). Accordingly, in *Drosophila melanogaster*, these two proteins compete for iron in the intestine and also have an inverted balance[100]. The expression of both encoding genes was significantly different in symbiotic and aposymbiotic weevils: *tsf-1* had an overall higher expression in symbiotic insects, peaking at D1 and then again D7/D9, while *fth-1* had a higher expression in aposymbiotic insects (peaking at D3, Figure 6). In *Asobara tabida*, a parasitoid wasp in the family Braconidae, Kremer et al. (2009)[101] also showed that both heavy and light chains of ferritin were over-expressed in aposymbiotic insects. Transferrin, in turn, might be necessary in symbiotic insects either to sequester iron and limit its availability as a means to control bacterial growth, or to directly deliver iron to the endosymbiont. The first hypothesis has been proposed as a response mechanism to pathogenic bacteria in *D. melanogaster*[102], while the latter was shown to take part in the homeostasis between *D. melanogaster* and *Spiroplasma poulsonii*[103]. On the other hand, aposymbiotic insects would need more ferritin to deal with iron excess (especially after emergence), or to transport it to other tissues. Moreover, the balance between both genes is important for the regulation of ferroptosis, an iron-dependent non-apoptotic cell death[104].

**Figure 6.**
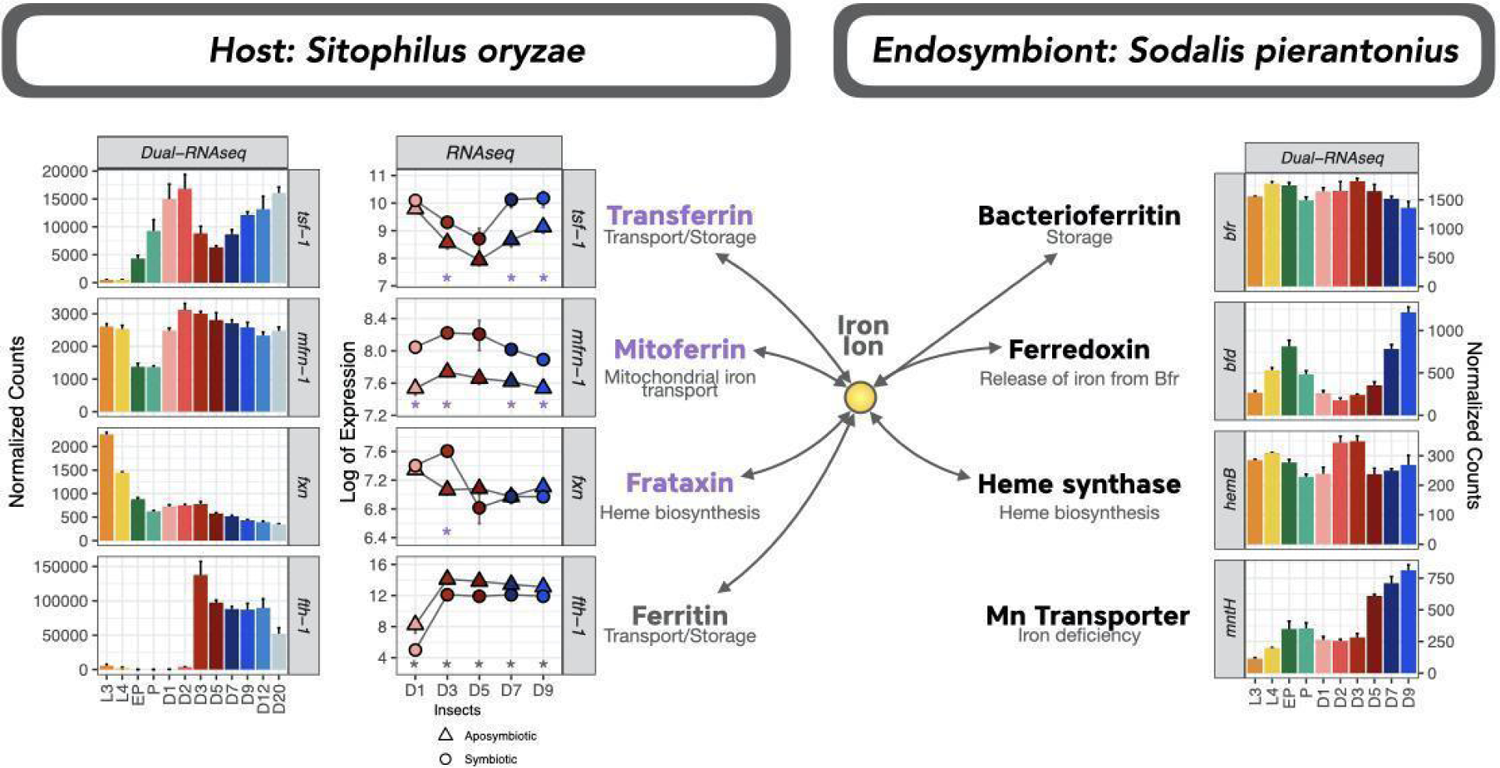
Expression of selected genes from iron metabolism in *S. oryzae* and *S. pierantonius* throughout the life cycle of the cereal weevil (dual RNA-seq) and/or differentially expressed between symbiotic and aposymbiotic insects (RNA-seq). Stars indicate in which stages we detected a significant up-regulation (purple) or down-regulation (gray) in symbiotic insects when compared to aposymbiotic within the RNA-seq dataset.

We found a coordinated bacterial gene expression of iron-related proteins: a bacterioferritin-associated ferredoxin (*bfd*), with a profile similar to *tsf-1* (belonging to supercluster SP19), and a bacterioferritin (*bfrA*) with a slight peak around D3 (supercluster SP17), but globally highly expressed throughout the weevil’s life cycle. In *Pseudomonas aeruginosa*, iron starvation downregulates *bfrA* while strongly upregulates *bfd* and vice-versa [105–107]. Thus, the coordinated expression of such proteins in *S. pierantonius* indicates that at D2/D3 iron is highly available for the bacteria, which drastically decreases until D9. This is particularly supported by the expression of several heme biosynthetic genes, which peak from D1 to D3/D5 (Supercluster SP16, Figure 2; *hemB* in Figure 6). *S. pierantonius* might be able to sense this decrease in iron content from D5 onwards and have different ways of coping with this (Supplementary Text S2B, refs[108,109]). Alternatively, this shortage of iron ions along with the demand for heme could be an outcome of the increased metabolic and respiratory activities during the symbiont growth period.

### Host recycling genes and bacterial stress response anticipate active bacterial clearance

We have previously shown that endosymbiont clearance and recycling involves bacteriocyte autophagy and apoptosis[40]. Even though the transcription of the autophagy factor *atg6* has been detected prior to active bacterial recycling[90], our initial hypothesis was that the accumulation of DOPA at D6 triggered apoptosis. More recently, we pinpointed that DOPA increase was concomitant with the endosymbiont clearance rather than anticipating it[41]. In accordance with Dell’Aglio et al. (2023)[41], we detected here a strong enrichment of autophagy and endosomal transport starting at D5 (supercluster SO70, Figure 2), before the active endosymbiont clearance. Moreover, all superclusters presenting a functional enrichment in autophagy (SO29, SO31, and SO39, Figure 2) had the lowest expression at D1 and D2, indicating the inhibition of autophagic genes at these early stages, and a gradual increase starting at D3. Finally, other cell-death and immune-related genes, belonging to enriched functions such as exocytosis, immune effector process and response to bacterium increase gradually from D5 to D20 (superclusters SO39 and SO69 from Figure 2). One example is the antimicrobial peptide Acanthoscurrin-2 (*acn-2*, supercluster SO69), which showed a prominent increase in expression of around 200-fold between D1 and D20.

The enrichment of endocytosis and autophagy at D5 and exocytosis and immune effectors (specified with the GO term antigen processing and presentation) at D5, D7, and D9 were further confirmed as symbiotic-specific processes (Superclusters S2 and S6, Figure 4). Hydrolase activity, essential for the final recycling of bacterial cell debris, is enriched specifically in symbiotic weevils at D9 (Supercluster S5). Nevertheless, the complex equilibrium between pro-apoptotic and pro-survival factors varies throughout the life cycle of the cereal weevil. Thus, the complex and symbiotic-specific expression of the Hippo signaling pathway, along with components from the Wnt pathway, possibly play a role in the regulation of programmed cell death of bacteriocytes (Figure 7).

**Figure 7.**
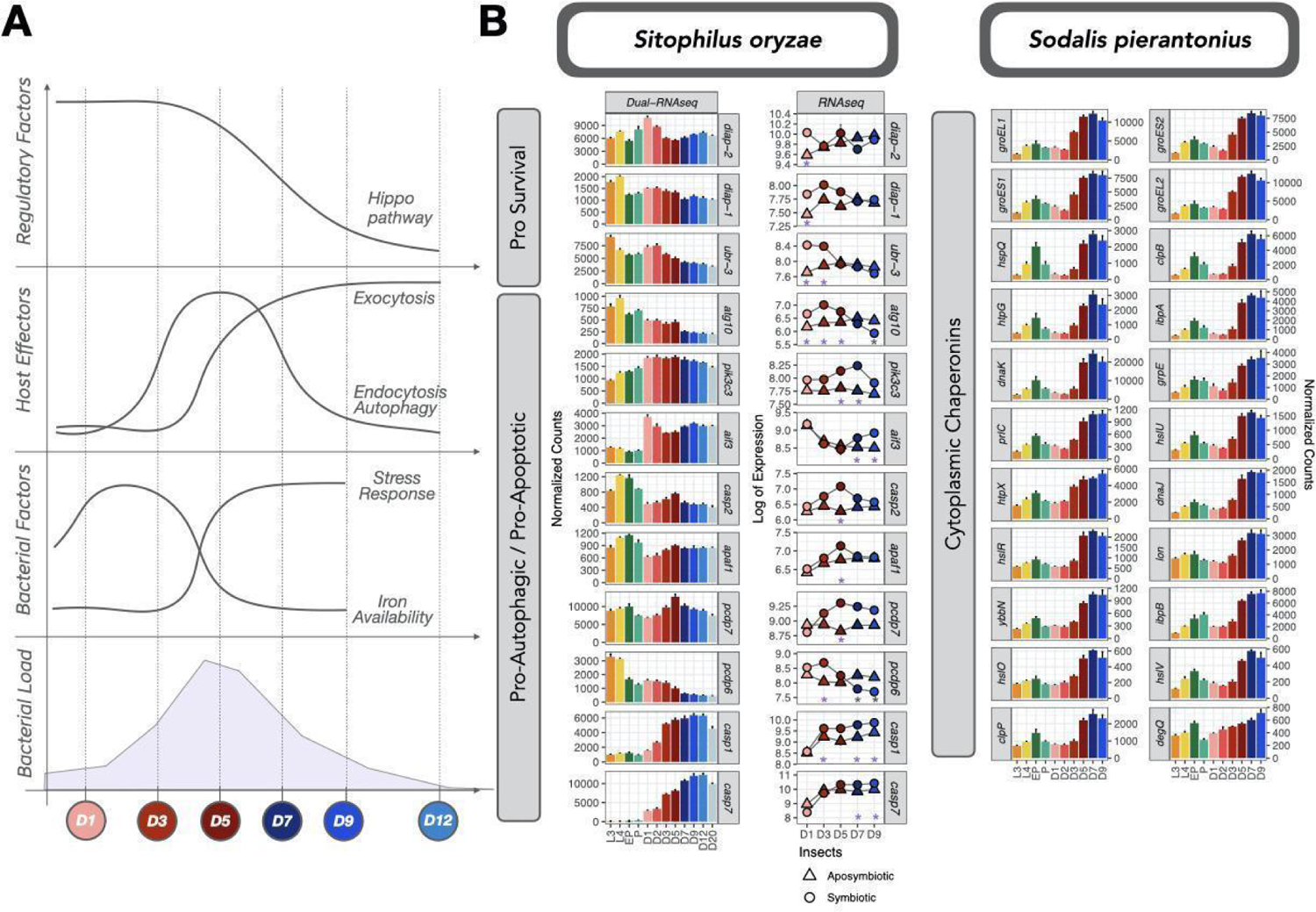
Insect and bacterial factors involved in the clearance of bacteria. (A) Schematic profiles of expression of host regulatory pathway Hippo, host effectors involved in exocytosis, endocytosis and autophagy as well as bacterial stress response genes depict a complex but coordinated regulation between both symbiotic partners. We also included the bacterial perceived iron availability (as predicted by the expression of iron sensing genes from Figure 6). (B) Selection of deregulated pro-survival pro-apoptotic / pro-autophagic genes from the cereal weevil as well as all cytoplasmic chaperonins involved in stress response from the endosymbiont throughout the life cycle of the cereal weevil (dual RNA-seq) and/or differentially expressed between symbiotic and aposymbiotic insects (RNA-seq). Stars indicate in which stages we detected a significant up-regulation (purple) or down-regulation (gray) in symbiotic insects when compared to aposymbiotic within the RNA-seq dataset.

While most effectors of cell death were induced in symbiotic insects at D5 and later, some presented a significant up-regulation at D3, such as *atg-10* and *pdcd-6*. Pro-survival genes *diap-1*, *diap-2* and *ubr-3* were highly expressed at the initial stages of adulthood and were downregulated at D5. The protein Ubr3 is known to regulate the activity of Diap1 in *Drosophila*[110], and its knockout was shown to induce caspase-dependent apoptosis. Accordingly, executioner caspases *casp-1* and *casp-2* were upregulated in symbiotic insects at D5. Even though no initiator caspase has been identified in the cereal weevil so far, the gene responsible for the activation of such a caspase was also highly induced at D5 (*apaf-1*).

In sum, apoptosis is likely functional as early as D5, with the induction of pro-apoptotic and pro-autophagic genes and depletion of pro-survival ones. At D3, even though the expression of some genes involved in programmed cell-death and autophagy is observed, pro-survival effectors most likely balance out the apoptotic effects, in accordance with our previous studies that show no distinct sign of apoptosis of bacteriocytes at D3[40].

Finally, we identified a major increase in expression of cytoplasmic chaperonins in the bacteria at D5, D7 and D9 (Figure 7, superclusters SP4, SP19 and SP23 from Figure 2, SP6, SP17 from Supplementary Figure S4). This enrichment in stress response genes is indeed the most significant enrichment within all bacterial clusters. Most chaperonins also have smaller peaks of expression at EP stage (Figure 7), during the period that endosymbionts exit bacteriomes and infect stem cells during metamorphosis, and thus are no longer protected from the immune system within bacteriocytes. In line with the fact that endocytosis, apoptosis, autophagy and other stress signals are activated prior to the active recycling of bacteria, some of these chaperones were already induced at D3, including the chaperonins *groEL* and *groES*. In weevils, GroEL has been previously described to have a central role in the inhibition of *S. pierantonius* division, through the interaction with ColA peptide[60]. The gene *ychH*, which codes for a putative inner membrane stress-induced protein, presented a similar profile. Interestingly, this gene was detected as the most repressed gene of *D. dadantii* during *Acyrthosiphon pisum* infection[61]. Finally, despite this widespread activation of stress genes, the general sigma factor responsible for the stress response *rpoH* did not present the same profile (supercluster SP2) (Supplementary Text S2C, refs[111–115]). Finally, by D5 most tRNAs are down-regulated (Supplementary Figure S9), indicating a general and abrupt translational arrest.

## Conclusions

The trade-off between host benefit and endosymbiont burden is central to the understanding of symbiosis establishment and maintenance. Whereas endosymbiont dynamics and load changes have been reported in several model systems, very little is known on genes and cellular pathways associated with host-symbiont trade-offs. In this study, we performed a comprehensive transcriptomic approach that includes dual RNA-seq and RNA-seq on symbiotic and aposymbiotic insects, to identify key functions in both insect and bacteria acting on bacterial dynamics at key development stages of cereals weevils.

We showed that at the larval stages, both bacteria and host activate cellular and metabolic processes, which may help anticipate the morphological changes occurring during metamorphosis, and store energy reserves before the fasting period begins. Moreover, we pinpointed a novel putative intracellular AMP, coding for a partial Diptericin-like (*dpt-like partial*), whose expression was highly correlated with bacterial load in symbiotic insects (Figure 3), likely involved in endosymbiont regulation.

The differential expression between symbiotic and aposymbiotic adults showed that endosymbiont presence greatly affects the gene expression within weevils. Regarding the contrasted bacterial dynamics, we examined the regulatory mechanisms involved in endosymbiont proliferation, bacterial load climax and recycling. Symbiotic adults showed an enrichment in functions related to carbohydrate metabolism, in agreement with our previous work demonstrating that endosymbiont proliferation relies solely on the availability of carbohydrates from diet intake[41]. Nevertheless, we highlighted biological functions intertwined and coregulated, including the Hippo signaling pathway, likely affecting bacterial dynamics, and bacterial cellular stress at the onset of endosymbiont recycling. Nutritional immunity through iron availability has been linked with both the Hippo signaling and apoptosis in *Drosophila*[116,117]. Moreover, the Hippo and Wnt signaling were shown to be important in the proliferative states of aphids[28,89]. Therefore, it is plausible that these biological functions could be central to the control of *S. pierantonius* load climax.

Even though early signals of apoptosis and autophagy were detected early on (D3), apoptosis is molecularly functional from D5 onwards, with both the induction of pro-apoptotic and pro-autophagic genes and the depletion of pro-survival ones from both bacteria and insect (Supplementary Text S2D, refs[68,118]). As the molecular recycling process begins, the bacteria sense this stress and activate a global stress response along with a translational arrest from D5 to D9. Following the bacterial recycling at D12/D20, we no longer detect changes in expression within the insect, which suggests that cell homeostasis within the gut is restored.

Future perspectives of this work include the integration with a reconstructed genome-wide metabolic network from both insect and bacterium, in order to gain insights into complete metabolic pathways. The complete study of these tightly regulated metabolic functions, which are at the center of symbiotic exchanges, will help to understand how the host and bacteria finely tune their gene expression and respond to physiological challenges constrained by insect development in a nutritionally limited ecological niche.

## Ethics approval and Consent to participate

Not applicable

## Consent for publication

All authors read and approved the final manuscript.

## Availability of data and materials

Raw sequencing data from this study have been deposited at the National Center for Biotechnology Information (NCBI) Sequence Read Archive (SRA) under PRJNA918957.

## Authors’ Contributions

AH, NP and RR conceived the original project. AV was responsible for all the molecular biology methods, with the help of EDA and OH. AV in collaboration with SH and BG constructed the dual RNA-seq libraries and produced the sequencing reads. AV constructed the RNA-seq libraries and produced the sequencing reads. MGF was responsible for the bioinformatic analyses of the dual RNA-seq and RNA-seq. MGF analyzed the data with the help of NP, RR, GC, and AH. MGF, AH, NP and RR wrote the manuscript with the help of CVM, GC and AZR.

## Acknowledgments

This work was funded by the ANR GREEN (ANR-17-CE20-0031-01–AH) and ANR UNLEASH (ANR UNLEASH-CE20-0015-01–RR). This work was performed using the computing facilities of the CC LBBE/PRABI. Dual RNA-seq library construction and sequencing was performed at the sequencing platform of the Institut de Génomique Fonctionnelle de Lyon, Ecole Normale Supérieure de Lyon, France. Sequencing for the RNA-seq library was performed by the GenomEast platform, a member of the ‘France Génomique’ consortium (ANR-10-INBS-0009). The authors wish to thank the DTAMB platform and Jihane Mennana for technical help regarding quantitative PCRs.

## Supplemental information

### Text

S1: Dual transcriptomic results

S2: Additional remarks on bacterial genes and enriched functions S3: Additional remarks on insect genes and enriched functions

S4: Metabolites predicted to be exchanged between insect and bacteria

### Figures#

S1: Bioinformatics methodology for dual RNA-seq

S2: Bacterial reads from dual RNA-seq

S3: Profile of dual RNA-seq superclusters from Sitophilus oryzae

S4: Profile of dual RNA-seq superclusters from Sodalis pierantonius

S5: Detailed analysis of putative deubiquitinase SseL

S6: Expression profiles of selected antimicrobial peptides from Sitophilus oryzae

S7: Profile of RNA-seq superclusters from symbiotic Sitophilus oryzae

S8: Profile of RNA-seq superclusters from aposymbiotic Sitophilus oryzae

S9: tRNA expression profiles from Sodalis pierantonius

### Tables

S1: Mapping statistics

S2: Dual RNA-seq gene counts from Sitophilus oryzae

S3: Dual RNA-seq gene counts from Sodalis pierantonius

S4: Dual RNA-seq DEGs from Sitophilus oryzae

S5: Dual RNA-seq DEGs from Sodalis pierantonius

S6: Dual RNA-seq LRT results from Sitophilus oryzae

S7: Dual RNA-seq LRT results from Sodalis pierantonius

S8: RNA-seq gene counts from symbiotic and aposymbiotic Sitophilus oryzae

S9: RNA-seq DEGs from symbiotic and aposymbiotic Sitophilus oryzae

S10: Profile of dual RNA-seq superclusters from Sitophilus oryzae

S11: Profile of dual RNA-seq superclusters from Sodalis pierantonius

S12: Profile of RNA-seq superclusters from symbiotic and aposymbiotic Sitophilus oryzae

S13: Dual RNA-seq functional enrichment analysis of supercluster from Sitophilus oryzae

S14: Dual RNA-seq functional enrichment analysis of supercluster from Sodalis pierantonius

S15: RNA-seq functional enrichment analysis of supercluster from symbiotic and aposymbiotic Sitophilus oryzae

S16: List of genes and abbreviations within main text

S17: List of signaling and immunity genes

