## Supplementary Figures for "Coordination of host and endosymbiont gene expression governs endosymbiont growth and elimination in the cereal weevil *Sitophilus* spp"

S1: Bioinformatics methodology for dual RNA-seq

S2: Euclidean clustering of sequencing samples

S3: PCA Clustering of sequencing samples

S4: Bacterial reads from dual RNA-seq

S5: Profile of dual RNA-seq superclusters from *Sitophilus oryzae*

S6: Profile of dual RNA-seq superclusters from *Sodalis pierantonius*

S7: Detailed analysis of putative deubiquitinase SseL

S8: Expression profiles of selected antimicrobial peptides from *Sitophilus oryzae*

S9: Profile of RNA-seq superclusters from symbiotic *Sitophilus oryzae*

S10: Profile of RNA-seq superclusters from aposymbiotic *Sitophilus oryzae*

S11: tRNA expression profiles from *Sodalis pierantonius*

S1

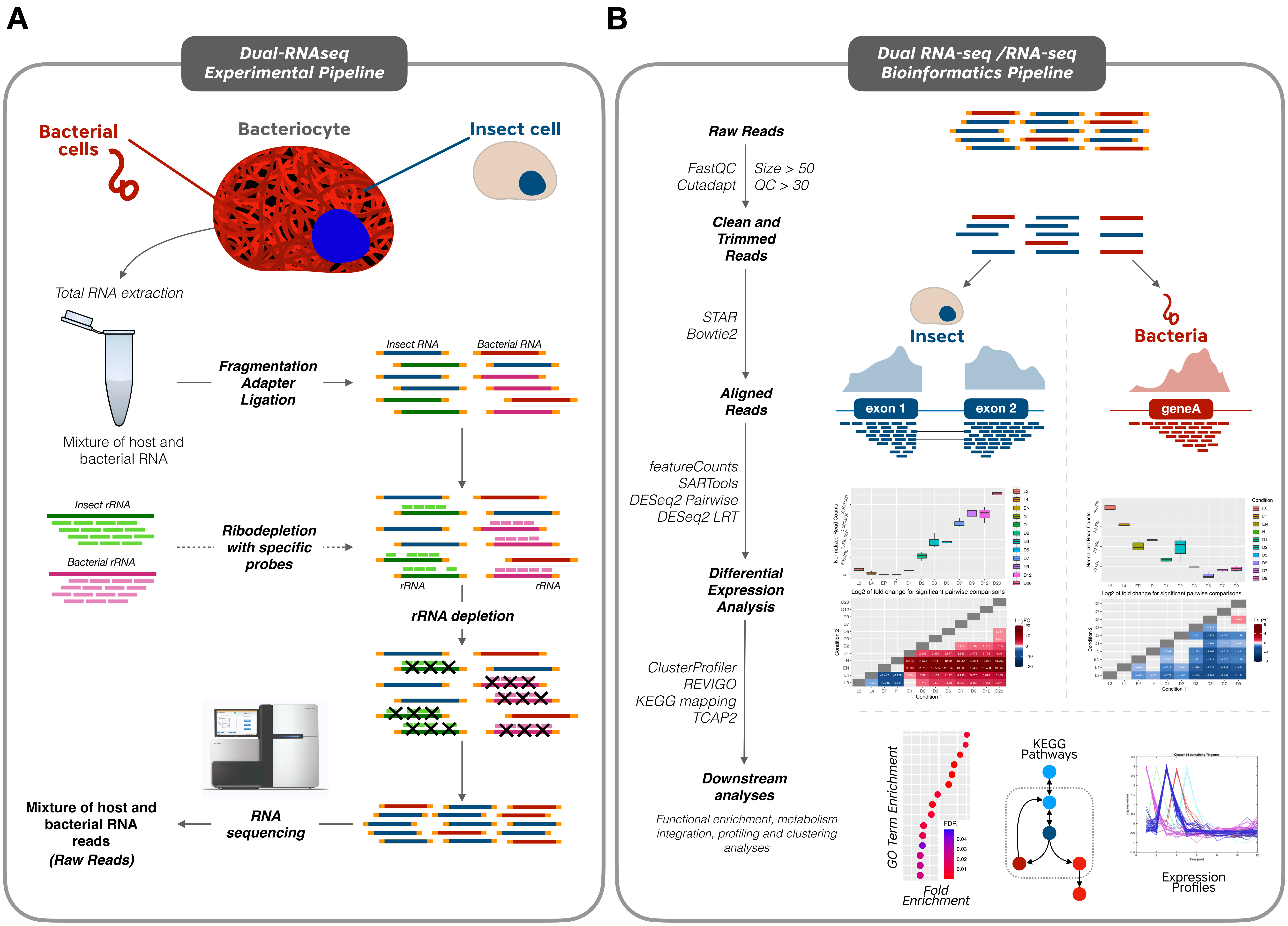

A
Dual-RNAseq Experimental Pipeline
Bacterial cells
Bacteriocyte
Insect cell
Total RNA extraction
Mixture of host and bacterial RNA
Fragmentation Adapter Ligation
Insect RNA
Bacterial RNA
Insect rRNA
Bacterial rRNA
Ribodepletion with specific probes
rRNA
rRNA
rRNA depletion
Mixture of host and bacterial RNA reads (Raw Reads)
RNA sequencing
B
Dual RNA-seq /RNA-seq Bioinformatics Pipeline
Raw Reads
FastQC
Cutadapt
Size > 50
QC > 30
Clean and Trimmed Reads
STAR
Bowtie2
Insect
Bacteria
Aligned Reads
exon 1
exon 2
geneA
featureCounts
SARTools
DESeq2 Pairwise
DESeq2 LRT
Differential Expression Analysis
Log2 of fold change for significant pairwise comparisons
Log2 of fold change for significant pairwise comparisons
ClusterProfiler
REVIGO
KEGG mapping
TCAP2
Downstream analyses
Functional enrichment, metabolism integration, profiling and clustering analyses
GO Term Enrichment
Fold Enrichment
FDR
KEGG Pathways
Expression Profiles

**S2**

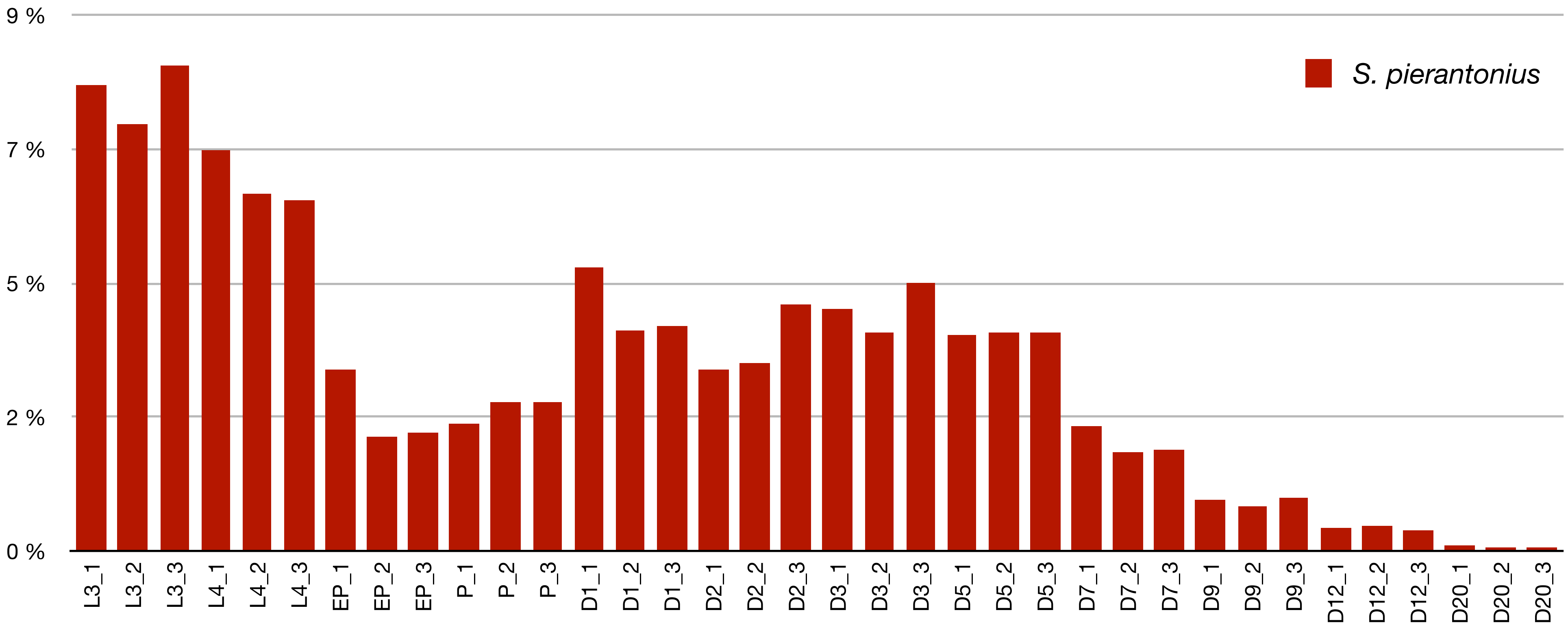
9 %
7 %
5 %
2 %
0 %
S. pierantonius
L3_1
L3_2
L3_3
L4_1
L4_2
L4_3
EP_1
EP_2
EP_3
P_1
P_2
P_3
D1_1
D1_2
D1_3
D2_1
D2_2
D2_3
D3_1
D3_2
D3_3
D5_1
D5_2
D5_3
D7_1
D7_2
D7_3
D9_1
D9_2
D9_3
D12_1
D12_2
D12_3
D20_1
D20_2
D20_3

# S3

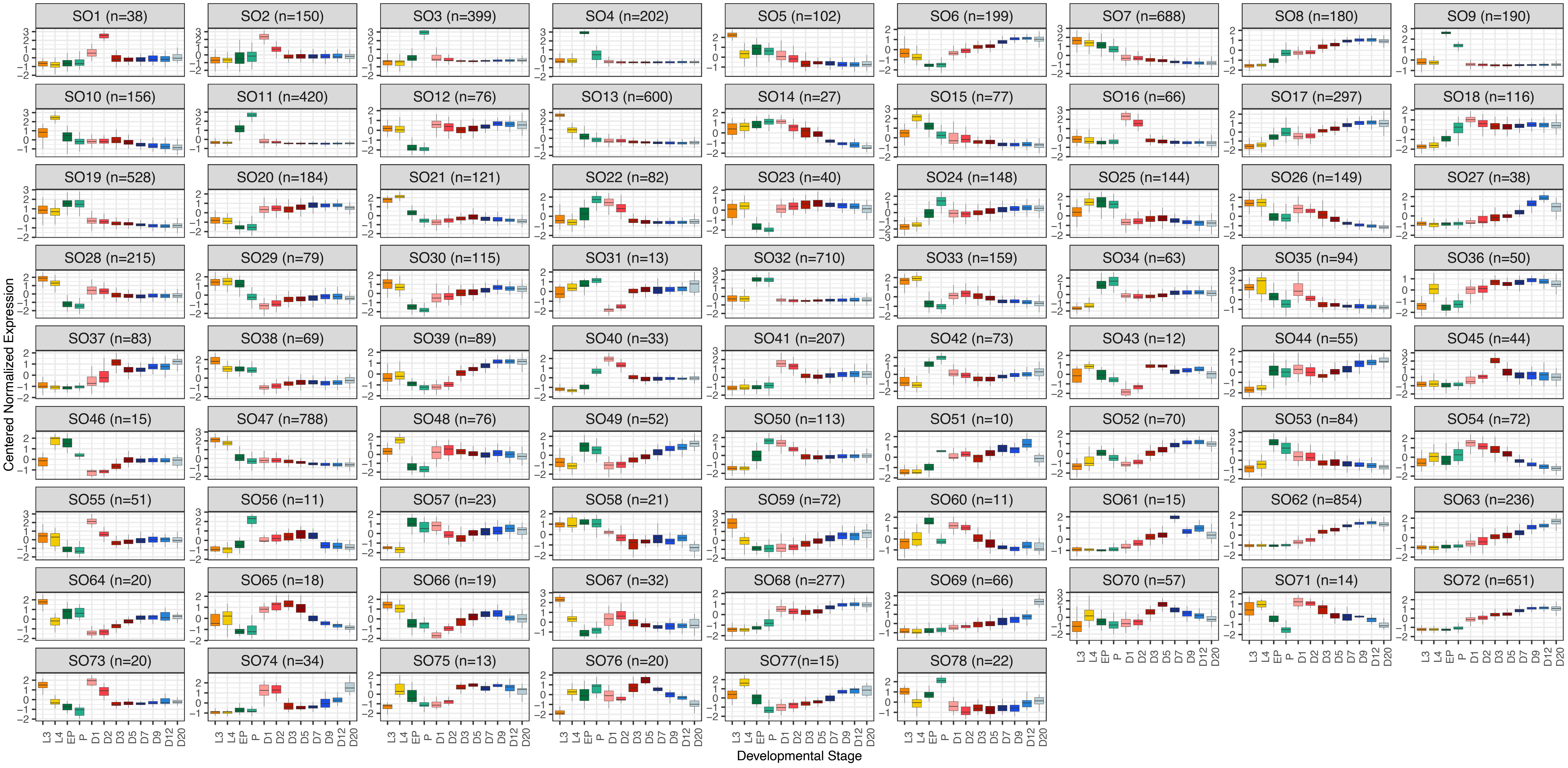
SO1 (n=38)
SO2 (n=150)
SO3 (n=399)
SO4 (n=202)
SO5 (n=102)
SO6 (n=199)
SO7 (n=688)
SO8 (n=180)
SO9 (n=190)
SO10 (n=156)
SO11 (n=420)
SO12 (n=76)
SO13 (n=600)
SO14 (n=27)
SO15 (n=77)
SO16 (n=66)
SO17 (n=297)
SO18 (n=116)
SO19 (n=528)
SO20 (n=184)
SO21 (n=121)
SO22 (n=82)
SO23 (n=40)
SO24 (n=148)
SO25 (n=144)
SO26 (n=149)
SO27 (n=38)
SO28 (n=215)
SO29 (n=79)
SO30 (n=115)
SO31 (n=13)
SO32 (n=710)
SO33 (n=159)
SO34 (n=63)
SO35 (n=94)
SO36 (n=50)
SO37 (n=83)
SO38 (n=69)
SO39 (n=89)
SO40 (n=33)
SO41 (n=207)
SO42 (n=73)
SO43 (n=12)
SO44 (n=55)
SO45 (n=44)
SO46 (n=15)
SO47 (n=788)
SO48 (n=76)
SO49 (n=52)
SO50 (n=113)
SO51 (n=10)
SO52 (n=70)
SO53 (n=84)
SO54 (n=72)
SO55 (n=51)
SO56 (n=11)
SO57 (n=23)
SO58 (n=21)
SO59 (n=72)
SO60 (n=11)
SO61 (n=15)
SO62 (n=854)
SO63 (n=236)
SO64 (n=20)
SO65 (n=18)
SO66 (n=19)
SO67 (n=32)
SO68 (n=277)
SO69 (n=66)
SO70 (n=57)
SO71 (n=14)
SO72 (n=651)
SO73 (n=20)
SO74 (n=34)
SO75 (n=13)
SO76 (n=20)
SO77(n=15)
SO78 (n=22)
Centered Normalized Expression
Developmental Stage
L3 L4 EP P D1 D2 D3 D5 D7 D9 D12 D20

# S4

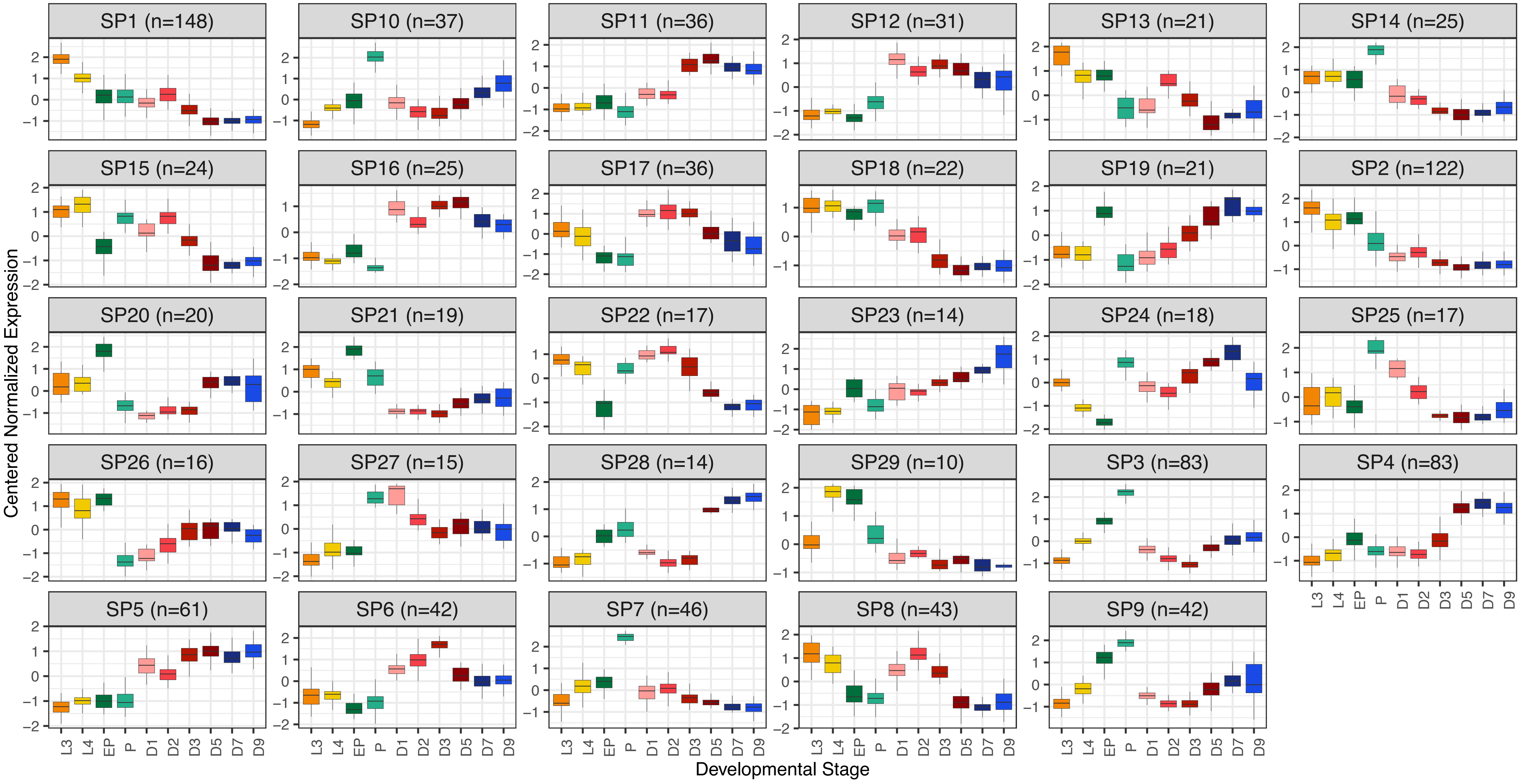
SP1 (n=148)
SP10 (n=37)
SP11 (n=36)
SP12 (n=31)
SP13 (n=21)
SP14 (n=25)
SP15 (n=24)
SP16 (n=25)
SP17 (n=36)
SP18 (n=22)
SP19 (n=21)
SP2 (n=122)
SP20 (n=20)
SP21 (n=19)
SP22 (n=17)
SP23 (n=14)
SP24 (n=18)
SP25 (n=17)
SP26 (n=16)
SP27 (n=15)
SP28 (n=14)
SP29 (n=10)
SP3 (n=83)
SP4 (n=83)
SP5 (n=61)
SP6 (n=42)
SP7 (n=46)
SP8 (n=43)
SP9 (n=42)
Centered Normalized Expression
L3 L4 EP P D1 D2 D3 D5 D7 D9
Developmental Stage

# S5

## A

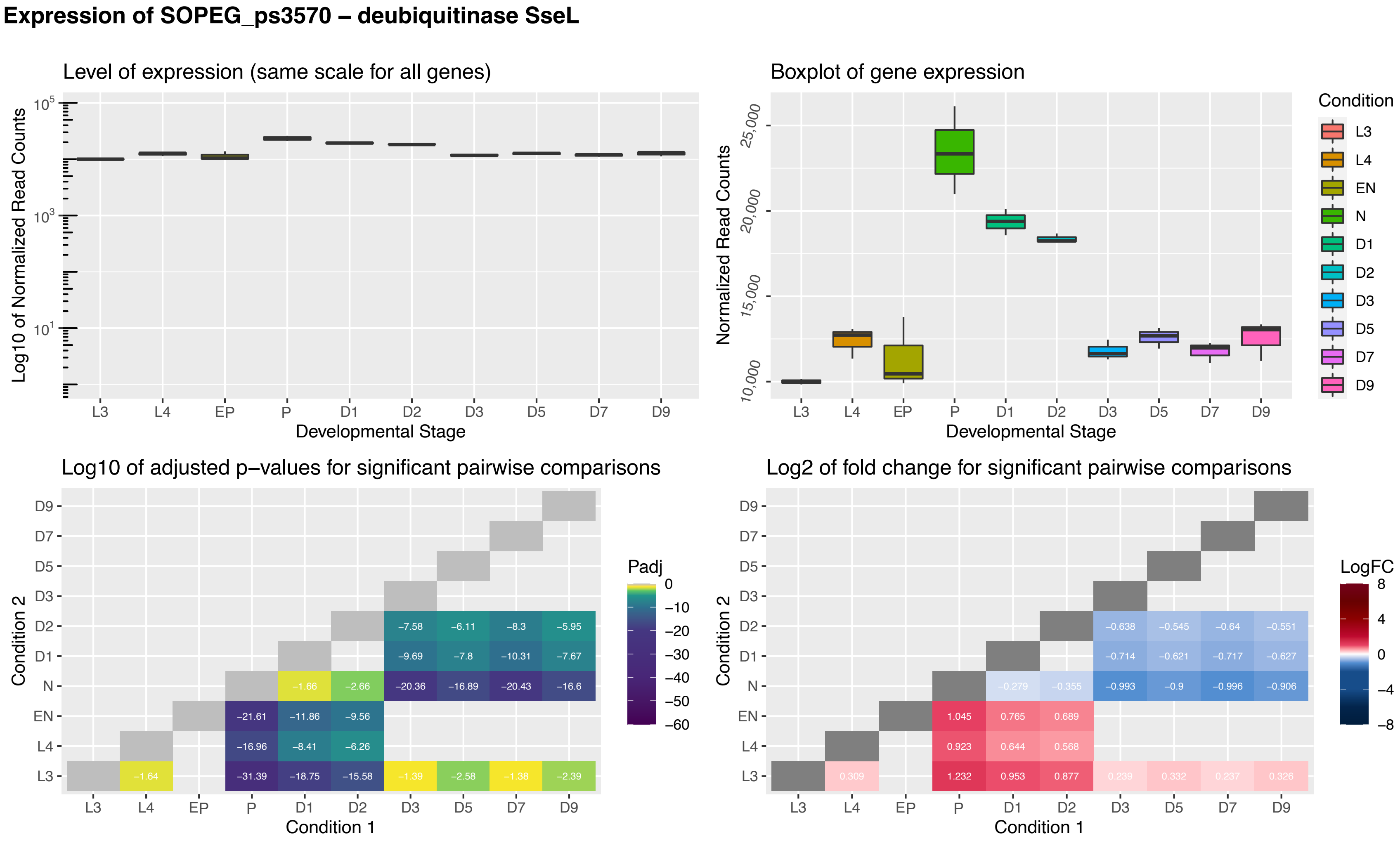


## B

>lcl|ORF1
MEYGYNFSSLETLKSAARNNDDFSIDALFTLACENSPVGLEAESFLFDLY
TGKEAGHPDLKEQLGKDSLKLCEIVQGRNKNKTPEEAWSIPDKVLIMAGF
ETQERSQQREEILEEINKNSKLLCLC

## C

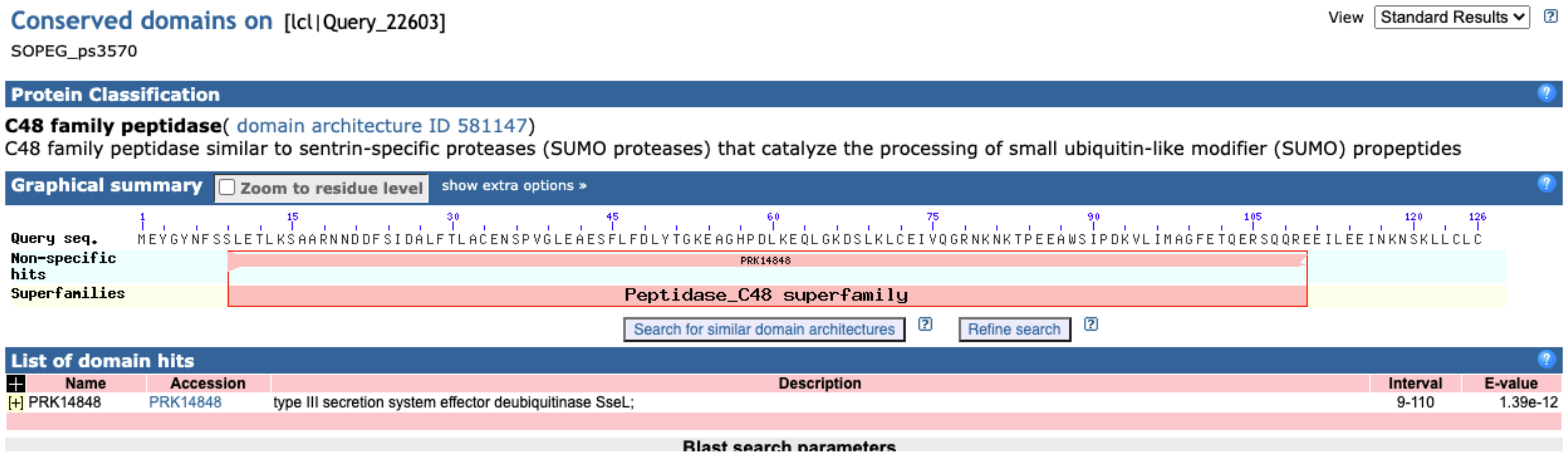

**S6**

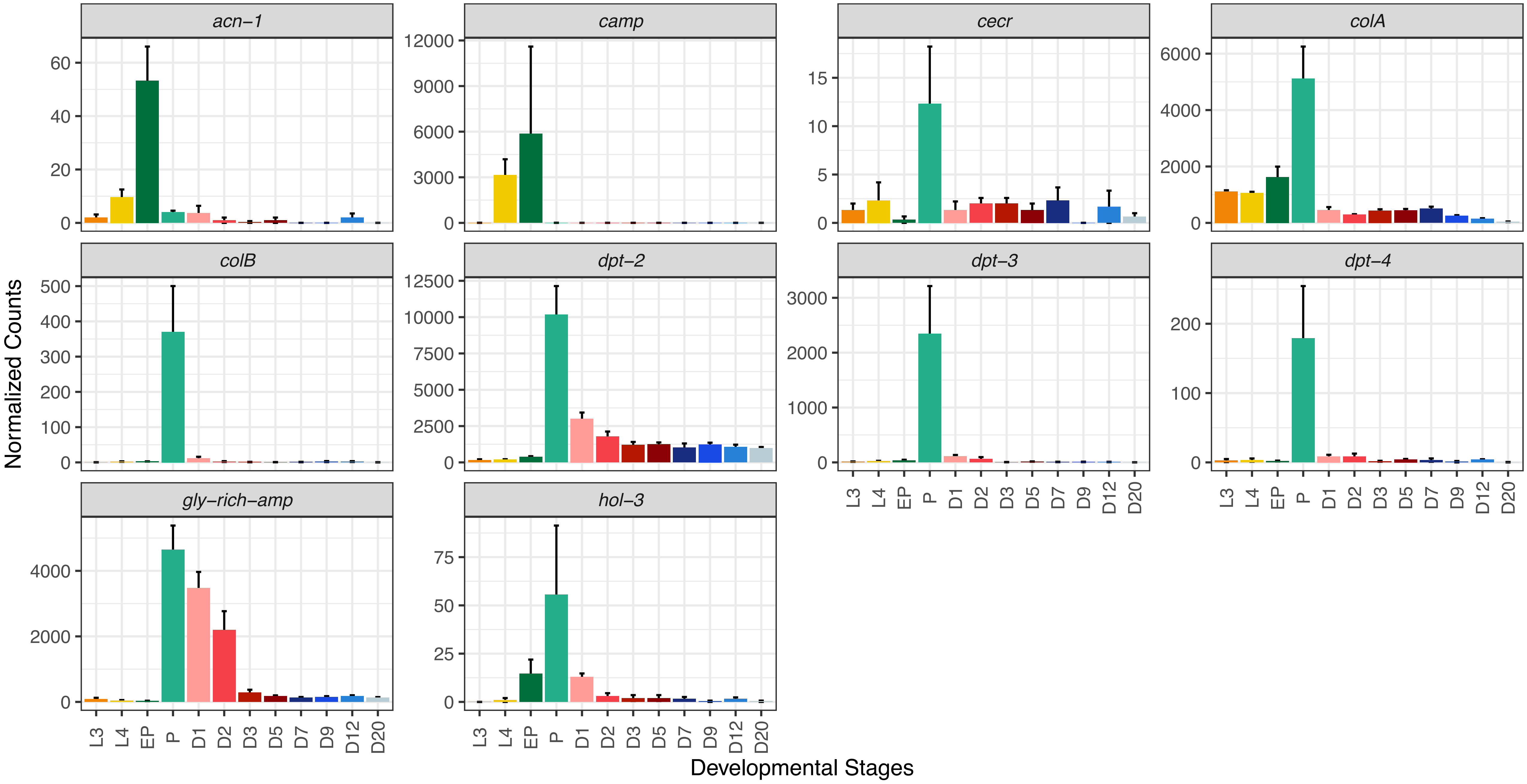

acn–1
camp
cecr
colA
colB
dpt–2
dpt–3
dpt–4
gly–rich–amp
hol–3
Normalized Counts
Developmental Stages
L3
L4
EP
P
D1
D2
D3
D5
D7
D9
D12
D20

**S7**

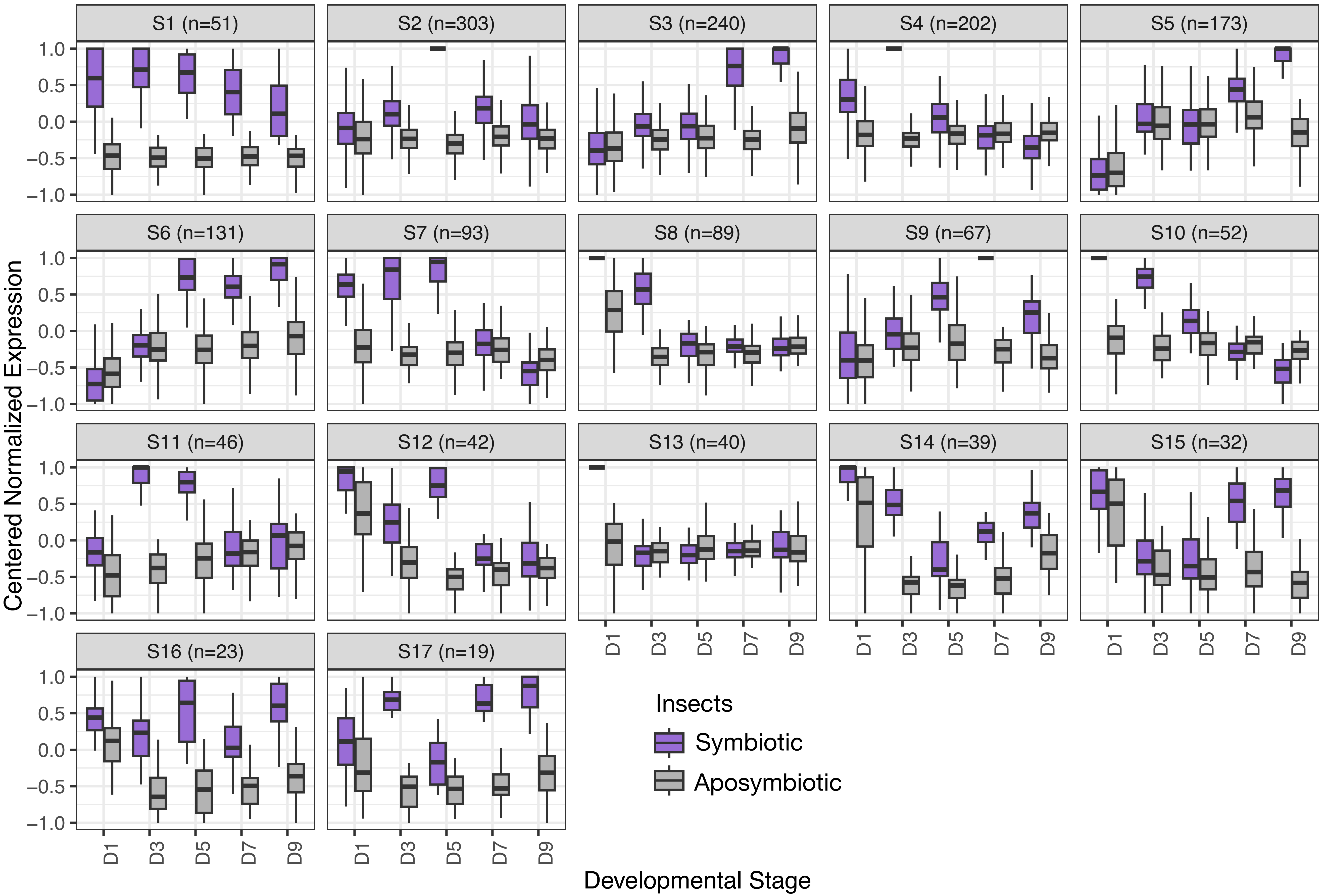
S1 (n=51)
S2 (n=303)
S3 (n=240)
S4 (n=202)
S5 (n=173)
S6 (n=131)
S7 (n=93)
S8 (n=89)
S9 (n=67)
S10 (n=52)
S11 (n=46)
S12 (n=42)
S13 (n=40)
S14 (n=39)
S15 (n=32)
S16 (n=23)
S17 (n=19)
1.0
0.5
0.0
−0.5
−1.0
D1
D3
D5
D7
D9
Centered Normalized Expression
Developmental Stage
Insects
Symbiotic
Aposymbiotic

**S8**

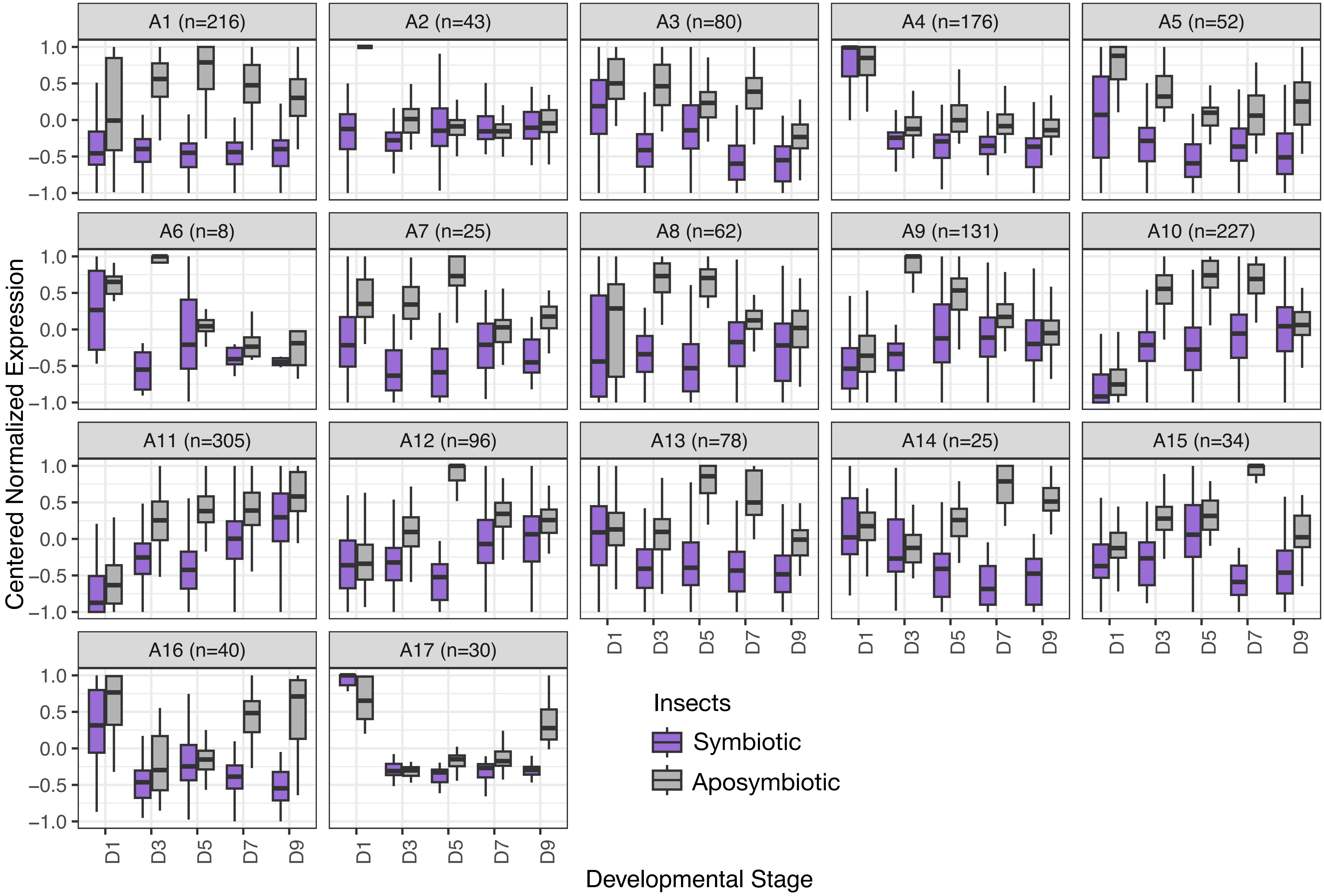

A1 (n=216)
A2 (n=43)
A3 (n=80)
A4 (n=176)
A5 (n=52)
A6 (n=8)
A7 (n=25)
A8 (n=62)
A9 (n=131)
A10 (n=227)
A11 (n=305)
A12 (n=96)
A13 (n=78)
A14 (n=25)
A15 (n=34)
A16 (n=40)
A17 (n=30)
1.0
0.5
0.0
−0.5
−1.0
Centered Normalized Expression
D1
D3
D5
D7
D9
Developmental Stage
Insects
Symbiotic
Aposymbiotic

**S9**

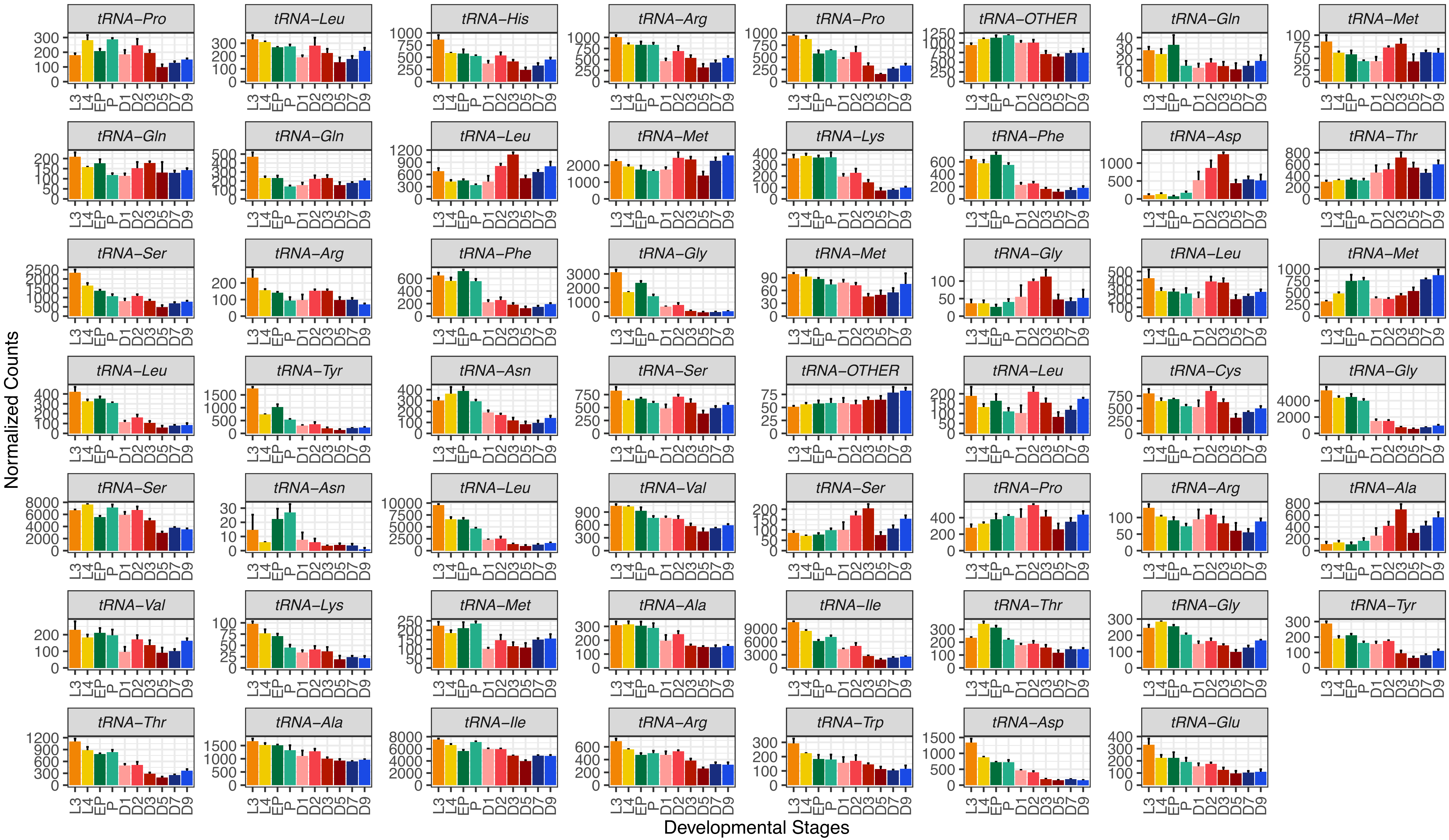
tRNA–Pro
tRNA–Leu
tRNA–His
tRNA–Arg
tRNA–Pro
tRNA–OTHER
tRNA–Gln
tRNA–Met
tRNA–Gln
tRNA–Gln
tRNA–Leu
tRNA–Met
tRNA–Lys
tRNA–Phe
tRNA–Asp
tRNA–Thr
tRNA–Ser
tRNA–Arg
tRNA–Phe
tRNA–Gly
tRNA–Met
tRNA–Gly
tRNA–Leu
tRNA–Met
tRNA–Leu
tRNA–Tyr
tRNA–Asn
tRNA–Ser
tRNA–OTHER
tRNA–Leu
tRNA–Cys
tRNA–Gly
tRNA–Ser
tRNA–Asn
tRNA–Leu
tRNA–Val
tRNA–Ser
tRNA–Pro
tRNA–Arg
tRNA–Ala
tRNA–Val
tRNA–Lys
tRNA–Met
tRNA–Ala
tRNA–Ile
tRNA–Thr
tRNA–Gly
tRNA–Tyr
tRNA–Thr
tRNA–Ala
tRNA–Ile
tRNA–Arg
tRNA–Trp
tRNA–Asp
tRNA–Glu
Normalized Counts
Developmental Stages
