## Supplementary Text for "Coordination of host and endosymbiont gene expression governs endosymbiont growth and elimination in the cereal weevil *Sitophilus* spp"

### Supplementary Text S1: Dual transcriptomic results

The dual RNA-seq recovered reads successfully from both insect and bacteria, ranging from 120 M to 200 M reads per sample (Supplementary Table S1). Depending on the tissue and time point, and in accordance with the bacterial load dynamics, the percentage of reads mapping to the bacteria varied from 8% of total trimmed reads (for L3 and L4) to less than 0.5% (~0.39% in D12 and ~0.07% in D20). The higher mapping of reads in L3 and L4 compared to later stages is explained mainly by the difference in tissue composition. While larval bacteriomes can be easily dissected from the fore-midgut junction, pupal and adult bacteriomes are tightly associated with the midgut and hence dual RNA-seq was conducted on dissected guts containing bacteriomes (Figure 1A). Focusing only on young adult bacterial reads, their proportion varies in accordance with the previously described bacterial load (Figure 1B and Supplementary Figure S2), denoting an overall correlation between bacterial load and bacterial gene expression. Due to the extremely low bacterial load at D12 vs D20, we were not able to perform differential expression analysis at these time points for *S. pierantonius*.

### Supplementary Text S2: Additional remarks on bacterial genes and enriched functions

**Supplementary Text S2A:** A complete bacterial operon (*arn*) acting on LPS modification through the addition of 4-amino-4-deoxy-L-arabinose to the outer membrane lipid A moiety and cationic antimicrobial peptide (CAMP) resistance was also induced at larval stages (supercluster SP1). This operon has previously been shown to be involved in increased antimicrobial peptide (AMP) resistance of the pathogenic free-living bacteria *Dickeya dadantii* (Costechareyre et al., 2013), when infecting the pea aphid *Acyrthosiphon pisum*. The *arn* operon, along with several other virulence response mechanisms, was described to be regulated by the PhoP/PhoQ proteins (Clayton et al., 2017). Here, genes encoding the PhoP/PhoQ system show higher expression at larval stages (*phoQ*, supercluster SP1, Figure 2 and *phoP*, supercluster SP2, Supplementary Figure S6), with the *phoQ* gene expression being the second most significant within the entire bacterial dataset (p-adj = 3.04E-266, Supplementary Table S7). At first sight, these results look intriguing when considering that endosymbionts are generally isolated from the host immune system and inhabit a stable environment where the requirement for sensory functions is expected to be relaxed or absent. Nevertheless, previous work on *Sodalis glossinidius*, the secondary endosymbiont of the tsetse fly which experiences both extra- and intracellular life-styles, showed that the PhoP/PhoQ system is required for the bacterial survival within the insect host by mediating the expression of genes involved in resistance to host AMPs (Pontes et al., 2011). This system was shown to be necessary for the transition from free-living to obligate endosymbiosis status in the tsetse fly (Clayton et al., 2017). It is important to notice that while the PhoP/PhoQ system could be essential in extracellular conditions in *S. glossinidius*, *S. pierantonius* is strictly intracellular during larval stages. Remarkably, all AMP encoding genes are downregulated within the weevil's bacteriocytes, with the notable exception of the *colA*, which continuously targets endosymbionts and inhibits their cell division (Login et al., 2011; Maire et al., 2018), and the *dpt-like partial*. The PhoP/PhoQ system activation in L3 and L4 may be connected to *colA* specific expression in bacteriomes.

**Supplementary Text S2B:** Knowing that different metals can occasionally substitute iron in activating at least some metalloenzymes, we looked at the expression of other metal-related genes, and found that the manganese transporter *mntH* (Figure 5), which can support the growth of iron-deprived bacteria(Makui et al., 2000), was highly upregulated from D5 to D9 (supercluster SP4). Interestingly, we also detected an increase in the expression of a Zinc uptake gene (*zntB*) from D1 to D3, which has been associated with nutritional immunity (Murdoch & Skaar, 2022).

**Supplementary Text S2C:** A transcriptional regulator involved in stress that follows the overall pattern of expression of the aforementioned chaperones (supercluster SP28 from Supplementary Figure S6), but which nonetheless has never been shown to regulate any of them, codes for the cell division protein BolA. The molecular mechanisms allowing a bacterial response to external signals are extremely complex and often involve two-component transduction systems (TCSs). Interestingly, *ompR*, a response regulator of the TCS pathway, which senses acid pH and osmotic stress, is also highly induced at D5 to D9 in *S. pierantonius* (supercluster SP4). This particular gene regulates the expression of virulence-related genes in *Salmonella* (Chakraborty et al., 2015; Lee et al., 2000)*,* *Yersinia* (Brzostek et al., 2007; Brzóstkowska et al., 2012), and *Shigella* (Bernardini et al., 1990). The bacteria could be presumably sensing an acidic pH within the host autophagic vesicles and by D5, most tRNAs are downregulated (Supplementary Figure S12), indicating a general and abrupt bacterial translational arrest.

**Supplementary Text S2D:** Ubiquitination-related processes, which have been extensively involved in immunity regulation and induction of both apoptosis and autophagy, especially in host-bacterial interactions (Vozandychova et al., 2021), were also enriched at D5 (both in the dual RNA-seq data, superclusters SO70 and SO76, and between symbiotic and aposymbiotic weevils, supercluster S2). As previously mentioned, the genome of *S. pierantonius* bears a single deubiquitinase *sseL*, which is known to inhibit autophagic clearance of cytoplasmic aggregates of *Salmonella* (Mesquita et al., 2012). This gene is induced at P, D1 and D2 stages in *S. pierantonius*, and this regulation decreases from D3 onwards (Supplementary Table S7). This suggests that, if the deubiquitinase SseL acts similarly in *S. pierantonius*, the bacteria could be actively helping the inhibition of apoptosis within the host at the early days of adulthood.

**Supplementary Text S3: Additional remarks on insect genes and enriched functions**

**Supplementary Text S3A:** We searched for insect genes that could directly impact bacterial gene expression during metamorphosis. This is the case of an insect putative Tol-Pal system protein (Figure 2, gene *tolA*), which is sharply induced (~30k fold expression) during the pupal stage (supercluster SO4). This gene is usually part of a known bacterial system involved in maintaining the integrity of the bacterial outer membrane (Yakhnina & Bernhardt, 2020) and to date there is no evidence for functional activity of this type of gene within any insect cell. We searched all RefSeq sequences from insects (taxid:6960) and from eukaryotes (taxid: 2759) and found only 7 species (from the order Coleoptera) bearing similar genes (Evalue < 10E-5, 50% identity; most of which are annotated as uncharacterized) (Table S3A.1). The bacteria itself has its own Tol-Pal system, and its profile precedes the insect's *tolA* expression: the bacterial *tolA* gene belongs to the supercluster SP2 (higher in larval stages).

**Table S3A.1: Hits against S. oryzae *tolA* from insects**

| Hit against XM_030897568.1 | Species | Query Cover | E-value | Per. Ident |
|---|---|---|---|---|
| XM_020018648.2 | Aethina tumida | 56% | 2.00E-24 | 66.05% |
| XM_020018754.1 | Aethina tumida | 25% | 1.00E-20 | 72.50% |
| XM_049961140.1 | Aethina tumida | 56% | 1.00E-19 | 65.05% |
| XM_049961145.1 | Aethina tumida | 43% | 2.00E-17 | 65.77% |
| XM_049961144.1 | Aethina tumida | 43% | 2.00E-17 | 65.77% |
| XM_049961143.1 | Aethina tumida | 43% | 2.00E-17 | 65.77% |
| XM_049961142.1 | Aethina tumida | 43% | 2.00E-17 | 65.77% |
| XM_049961141.1 | Aethina tumida | 43% | 2.00E-17 | 65.77% |
| XM_020017942.2 | Aethina tumida | 21% | 2.00E-17 | 70.32% |
| XM_020018792.2 | Aethina tumida | 42% | 7.00E-17 | 65.58% |
| XM_020018172.1 | Aethina tumida | 42% | 9.00E-16 | 65.37% |
| XM_020018183.2 | Aethina tumida | 43% | 9.00E-16 | 65.25% |
| XM_020017918.1 | Aethina tumida | 23% | 1.00E-13 | 68.91% |
| XM_018717288.1 | Anoplophora glabripennis | 55% | 1.00E-33 | 67.43% |
| XM_018717291.1 | Anoplophora glabripennis | 56% | 4.00E-26 | 65.97% |
| XM_018717292.1 | Anoplophora glabripennis | 55% | 6.00E-24 | 65.66% |
| XM_018717289.1 | Anoplophora glabripennis | 17% | 5.00E-19 | 73.45% |
| XM_018717295.1 | Anoplophora glabripennis | 33% | 1.00E-14 | 66.47% |
| XM_050450282.1 | Anthonomus grandis grandis | 58% | 1.00E-57 | 69.58% |
| XM_048670011.1 | Dendroctonus ponderosae | 76% | 3.00E-98 | 71.23% |
| XM_048670010.1 | Dendroctonus ponderosae | 76% | 3.00E-98 | 71.23% |
| XM_019909022.2 | Dendroctonus ponderosae | 76% | 3.00E-98 | 71.23% |
| XM_050643152.1 | Diabrotica virgifera virgifera | 59% | 4.00E-27 | 66.61% |
| XM_028283824.2 | Diabrotica virgifera virgifera | 18% | 1.00E-19 | 73.22% |
| XM_023165374.1 | Leptinotarsa decemlineata | 53% | 1.00E-26 | 66.48% |
| XM_023169520.1 | Leptinotarsa decemlineata | 62% | 3.00E-16 | 65.45% |
| XM_023162173.1 | Leptinotarsa decemlineata | 12% | 9.00E-16 | 76.98% |
| XM_023157170.1 | Leptinotarsa decemlineata | 21% | 1.00E-14 | 73.97% |
| XM_023157164.1 | Leptinotarsa decemlineata | 24% | 1.00E-14 | 71.59% |
| XM_008197753.2 | Tribolium castaneum | 17% | 2.00E-06 | 69.83% |
| XM_008197752.2 | Tribolium castaneum | 17% | 2.00E-06 | 69.83% |
| XM_044408620.1 | Tribolium madens | 42% | 5.00E-06 | 63.29% |

**Supplementary Text S3B:** Two examples of genes highly expressed in aposymbiotic insects in comparison to symbiotic ones presented in Figure 3 were *rutC*, an ubiquitous gene with diverse biological and metabolic functions (Niehaus et al., 2015), and *jagged*, a gene that interacts with the Notch protein to regulate cell-cell communication (Zeronian et al., 2021) and to determine cell-fate (Artavanis-Tsakonas et al., 1999). Moreover, when checking genes commonly induced in more than two stages (Superclusters A1, A9, A10 and A11, Supplementary Figure S11), we detected significant enrichments in peptidase activity, amino acid catabolism and small molecule catabolism after insect emergence. We also found several transmembrane transporters enriched in aposymbiotic insects, which could help to increase the uptake of essential metabolites, not readily available in aposymbiotic insects. In addition, Bonnot et al. (1998) showed that the feces of aposymbiotic weevils contained more residues than the feces of symbiotic ones, due to a lower conversion rate of metabolites within the gut of aposymbiotic insects, related to the fact that a great portion of what symbiotic insects uptake is redirected to the endosymbionts. Thus, the genes and functional enrichment induced in aposymbiotic young adults provided insights into specific biological functions necessary to deal with either the excess of metabolites that are not used by the bacteria or the deprivation of compounds produced by the bacteria, in insects lacking the endosymbiont.

**Supplementary Text S3C:** Ubiquitination, vacuolar transport and endocytosis are general processes involved in programmed cell death which were detected as differentially regulated between symbiotic and aposymbiotic weevils, particularly at D5 (Supercluster S2, Figure 4). Ubiquitin-related processes seem to be differentially regulated between symbiotic and aposymbiotic insects in different stages of the life-cycle of the cereal weevil, not only at D5. The gene coding for Ubiquitin domain-containing protein 2 (*ubtd-2*, Figure 3), for instance, was up-regulated at D7 and D9 in symbiotic insects. Ubiquitin-ligases are also the main effectors on the clearance of intestinal epithelium pathogens in the nematode *Caenorhabditis elegans*. The intracellular pathogen response (IPR) pathway is a physiological program that promotes resistance to proteotoxic stress, independent of other known stress response pathways (Reddy et al., 2017). Even though many of the effector genes within this pathway are specific to nematodes, the purine nucleoside phosphorylase (PNP), an enzyme involved in salvaging purines and which was described as a negative regulator of the IPR pathway, was found within our dataset as one of the genes most significantly differentially expressed between aposymbiotic and symbiotic insects (p-adj = 8.9E-110). This protein is thought to directly act in intestinal epithelial cells of *C. elegans* to regulate defense against viruses and microsporidia

(Tecle et al., 2021), and the infection with these intracellular pathogens was impaired by the loss of PNP. It is possible that similarly to what was proposed in *C. elegans*, perturbations in purine metabolism are used as cues in the *S. oryzae*'s gut epithelium to either prevent early recycling of bacteria or to actively promote the recycling period. This gene is highly expressed from larval stages up to the initial stages of adulthood and is sharply down-regulated from D5 onwards.

**Supplementary Text S3D:** In *Drosophila*, the intestinal epithelium turnover relies on the antagonistic activity of pro-apoptotic Notch and anti-apoptotic EGFR (Reiff et al., 2019). Notch is a cell-surface receptor that interacts with transmembrane ligands, leading to the cleavage of its intracellular domain (NICD) and migration to the nucleus, where it regulates transcription (Kopan, 2012). The *notch* gene had an increase in expression at D1 and D3 specifically in symbiotic insects (Supercluster S10), which gradually declined concomitantly to bacterial recycling. The two Presenilin (*psn*) coding genes, which were up-regulated at D5 in symbiotic insects (Supercluster S2), belong to the protease complex gamma secretase, the complex responsible for the cleavage of the NICD. Moreover, evidence suggests that other genes downstream the Notch cascade can affect other signaling pathways, such as Wnt and Hippo (Nagel et al., 2001; Watanabe et al., 2017).

**Supplementary Text S4: Metabolites predicted to be exchanged between insect and bacteria**

While weevils mainly feed on complex polysaccharides from cereal grains, such as starch and cellulose, symbiotic bacteria generally uptake derived carbohydrates, mostly converted to monosaccharides. We previously predicted which metabolites were exchanged during the initial stages of adulthood based on the complete genomes of both partners (Oakeson et al., 2014; Parisot et al., 2021), and pinpointed, in addition to monosaccharides, essential amino acids, cofactors and nucleotide precursors (Parisot et al., 2021). Host genes enriched at the initial stages of adulthood (D1 and D2), prior to insect emergence from the grain (superclusters SO1, SO2, SO40, SO41, from Figure 2; SO16, SO54, and SO55 from Supplementary Figure S5), had a significant functional enrichment related to ribosome biogenesis, rRNA processing, actin cytoskeleton, muscle system process, suggesting tissue development (especially at D1) before imminent feeding and emergence from the grain.

We recently showed that at D1, insects are motionless and unable to feed (Dell'Aglio et al., 2023), a process that restarts at D2. Our data point to the synthesis of novel proteins necessary for midgut food uptake and muscle peristaltic contraction. Other functions activated at these initial stages of adulthood are juvenile hormone catabolism, indicating the end of metamorphosis, and hydrolase activity, including glycosidases, a pectinesterase, endoglucanases and an exoglucanase B - *exgB*, showing that the degradation of cellulose and other insoluble plant cell wall sugars might be essential, in addition to starch inside the grains. Concomitantly and before emergence, a pectin degradation protein encoding gene is induced in bacteria at D1. Moreover, at the P stage, a complete pathway for the degradation of galacturonate - the pectin monomeric unit (*uxaABC*), along with two known regulators of these functions (*uxuR* and *kdgR*) in other bacteria (James & Hugouvieux-Cotte-Pattat, 1996; Nasser et al., 1994; Rodionov et al., 2000) also present an increased expression.

**Supplementary Text References**


Artavanis-Tsakonas, S., Rand, M. D., & Lake, R. J. (1999). Notch signaling: cell fate control and signal integration in development. *Science*, *284*(5415), 770–776.

Bernardini, M. L., Fontaine, A., & Sansonetti, P. J. (1990). The two-component regulatory system ompR-envZ controls the virulence of Shigella flexneri. *Journal of Bacteriology*, *172*(11). https://doi.org/10.1128/jb.172.11.6274-6281.1990

Brzostek, K., Brzóstkowska, M., Bukowska, I., Karwicka, E., & Raczkowska, A. (2007). OmpR negatively regulates expression of invasin in Yersinia enterocolitica. *Microbiology* , *153*(Pt 8). https://doi.org/10.1099/mic.0.2006/003202-0

Brzóstkowska, M., Raczkowska, A., & Brzostek, K. (2012). OmpR, a response regulator of the two-component signal transduction pathway, influences inv gene expression in Yersinia enterocolitica O9. *Frontiers in Cellular and Infection Microbiology*, *2*. https://doi.org/10.3389/fcimb.2012.00153

Chakraborty, S., Mizusaki, H., & Kenney, L. J. (2015). A FRET-based DNA biosensor tracks OmpR-dependent acidification of Salmonella during macrophage infection. *PLoS Biology*, *13*(4), e1002116.

Clayton, A. L., Enomoto, S., Su, Y., & Dale, C. (2017). The Regulation of Antimicrobial Peptide Resistance in the Transition to Insect Symbiosis. *Molecular Microbiology*, *103*(6), 958.

Costechareyre, D., Chich, J.-F., Strub, J.-M., Rahbé, Y., & Condemine, G. (2013). Transcriptome of Dickeya dadantii infecting Acyrthosiphon pisum reveals a strong defense against antimicrobial peptides. *PloS One*, *8*(1), e54118.

Dell'Aglio, E., Lacotte, V., Peignier, S., Rahioui, I., Benzaoui, F., Vallier, A., Da Silva, P., Desouhant, E., Heddi, A., & Rebollo, R. (2023). Weevil Carbohydrate Intake Triggers Endosymbiont Proliferation: A Trade-Off between Host Benefit and Endosymbiont Burden. *mBio*. https://doi.org/10.1128/mbio.03333-22

James, V., & Hugouvieux-Cotte-Pattat, N. (1996). Regulatory systems modulating the transcription of the pectinase genes of Erwinia chrysanthemi are conserved in Escherichia coli. *Microbiology* , *142 ( Pt 9)*. https://doi.org/10.1099/00221287-142-9-2613

Kopan, R. (2012). Notch signaling. *Cold Spring Harbor Perspectives in Biology*, *4*(10). https://doi.org/10.1101/cshperspect.a011213

Lee, A. K., Detweiler, C. S., & Falkow, S. (2000). OmpR Regulates the Two-Component System SsrA-SsrB in Salmonella Pathogenicity Island 2. *Journal of Bacteriology*, *182*(3), 771.

Login, F. H., Balmand, S., Vallier, A., Vincent-Monégat, C., Vigneron, A., Weiss-Gayet, M., Rochat, D., & Heddi, A. (2011). Antimicrobial peptides keep insect endosymbionts under control. *Science* , *334*(6054). https://doi.org/10.1126/science.1209728

Maire, J., Vincent-Monégat, C., Masson, F., Zaidman-Rémy, A., & Heddi, A. (2018). An IMD-like pathway mediates both endosymbiont control and host immunity in the cereal weevil Sitophilus spp. *Microbiome*, *6*(1). https://doi.org/10.1186/s40168-017-0397-9

Makui, H., Roig, E., Cole, S. T., Helmann, J. D., Gros, P., & Cellier, M. F. (2000). Identification of the Escherichia coli K-12 Nramp orthologue (MntH) as a selective divalent metal ion transporter. *Molecular Microbiology*, *35*(5). https://doi.org/10.1046/j.1365-2958.2000.01774.x

Mesquita, F. S., Thomas, M., Sachse, M., Santos, A. J. M., Figueira, R., & Holden, D. W. (2012). The Salmonella deubiquitinase SseL inhibits selective autophagy of cytosolic aggregates. *PLoS Pathogens*, *8*(6), e1002743.

Murdoch, C. C., & Skaar, E. P. (2022). Nutritional immunity: the battle for nutrient metals at the host-pathogen interface. *Nature Reviews. Microbiology*, *20*(11). https://doi.org/10.1038/s41579-022-00745-6

Nagel, A. C., Wech, I., & Preiss, A. (2001). Scalloped and strawberry notch are target genes of Notch signaling in the context of wing margin formation in Drosophila. *Mechanisms of Development*, *109*(2), 241–251.

Nasser, W., Reverchon, S., Condemine, G., & Robert-Baudouy, J. (1994). Specific interactions of Erwinia chrysanthemi KdgR repressor with different operators of genes involved in pectinolysis. *Journal of Molecular Biology*, *236*(2). https://doi.org/10.1006/jmbi.1994.1155

Niehaus, T. D., Gerdes, S., Hodge-Hanson, K., Zhukov, A., Cooper, A. J. L., ElBadawi-Sidhu, M., Fiehn, O., Downs, D. M., & Hanson, A. D. (2015). Genomic and experimental evidence for multiple metabolic functions in the RidA/YjgF/YER057c/UK114 (Rid) protein family. *BMC Genomics*, *16*(1), 382.

Oakeson, K. F., Gil, R., Clayton, A. L., Dunn, D. M., von Niederhausern, A. C., Hamil, C., Aoyagi, A., Duval, B., Baca, A., Silva, F. J., Vallier, A., Jackson, D. G., Latorre, A., Weiss, R. B., Heddi, A., Moya, A., & Dale, C. (2014). Genome degeneration and adaptation in a nascent stage of symbiosis. *Genome Biology and Evolution*, *6*(1). https://doi.org/10.1093/gbe/evt210

Parisot, N., Vargas-Chávez, C., Goubert, C., Baa-Puyoulet, P., Balmand, S., Beranger, L., Blanc, C., Bonnamour, A., Boulesteix, M., Burlet, N., Calevro, F., Callaerts, P., Chancy, T., Charles, H., Colella, S., Da Silva Barbosa, A., Dell'Aglio, E., Di Genova, A., Febvay, G., … Heddi, A. (2021). The transposable element-rich genome of the cereal pest Sitophilus oryzae. *BMC Biology*, *19*(1). https://doi.org/10.1186/s12915-021-01158-2

Pontes, M. H., Smith, K. L., De Vooght, L., Den Abbeele J, V., & Dale, C. (2011). Attenuation of the sensing capabilities of PhoQ in transition to obligate insect-bacterial association. *PLoS Genetics*, *7*(11). https://doi.org/10.1371/journal.pgen.1002349

Reddy, K. C., Dror, T., Sowa, J. N., Panek, J., Chen, K., Lim, E. S., Wang, D., & Troemel, E. R. (2017). An Intracellular Pathogen Response Pathway Promotes Proteostasis in C. elegans. *Current Biology: CB*, *27*(22), 3544–3553.e5.

Reiff, T., Antonello, Z. A., Ballesta-Illán, E., Mira, L., Sala, S., Navarro, M., Martinez, L. M., & Dominguez, M. (2019). Notch and EGFR regulate apoptosis in progenitor cells to ensure gut homeostasis in Drosophila. *The EMBO Journal*, *38*(21), e101346.

Rodionov, D. A., Mironov, A. A., Rakhmaninova, A. B., & Gelfand, M. S. (2000). Transcriptional regulation of transport and utilization systems for hexuronides, hexuronates and hexonates in gamma purple bacteria. *Molecular Microbiology*, *38*(4). https://doi.org/10.1046/j.1365-2958.2000.02115.x

Tecle, E., Chhan, C. B., Franklin, L., Underwood, R. S., Hanna-Rose, W., & Troemel, E. R. (2021). The purine nucleoside phosphorylase pnp-1 regulates epithelial cell resistance to infection in C. elegans. *PLoS Pathogens*, *17*(4), e1009350.

Vozandychova, V., Stojkova, P., Hercik, K., Rehulka, P., & Stulik, J. (2021). The Ubiquitination System within Bacterial Host–Pathogen Interactions. *Microorganisms*, *9*(3). https://doi.org/10.3390/microorganisms9030638

Watanabe, Y., Miyasaka, K. Y., Kubo, A., Kida, Y. S., Nakagawa, O., Hirate, Y., Sasaki, H., & Ogura, T. (2017). Notch and Hippo signaling converge on Strawberry Notch 1 (Sbno1) to synergistically activate Cdx2 during specification of the trophectoderm. *Scientific Reports*, *7*, 46135.

Yakhnina, A. A., & Bernhardt, T. G. (2020). The Tol-Pal system is required for peptidoglycan-cleaving enzymes to complete bacterial cell division. *Proceedings of the National Academy of Sciences of the United States of America*, *117*(12), 6777–6783.

Zeronian, M. R., Klykov, O., Portell I de Montserrat, J., Konijnenberg, M. J., Gaur, A., Scheltema, R. A., & Janssen, B. J. C. (2021). Notch-Jagged signaling complex defined by an interaction mosaic. *Proceedings of the National Academy of Sciences of the United States of America*, *118*(30). https://doi.org/10.1073/pnas.2102502118
